# TWIST1 expression is associated with high-risk Neuroblastoma and promotes Primary and Metastatic Tumor Growth

**DOI:** 10.1101/2021.03.22.435811

**Authors:** Maria-Vittoria Sepporta, Viviane Praz, Katia Balmas Bourloud, Jean-Marc Joseph, Nicolas Jauquier, Nicolo’ Riggi, Katya Nardou-Auderset, Audrey Petit, Jean-Yves Scoazec, Hervé Sartelet, Raffaele Renella, Annick Mühlethaler-Mottet

**Author notes:** **Competing interests**, The authors declare no competing interests.

## Abstract

The embryonic transcription factors TWIST1/2 are frequently overexpressed in cancer, acting as multifunctional oncogenes. Here we investigate their role in neuroblastoma (NB), a heterogeneous childhood malignancy ranging from spontaneous regression to dismal outcomes despite multimodal therapy. We first reveal the association of TWIST1 expression with poor survival and metastasis in primary NB, while TWIST2 correlates with good prognosis. Secondly, suppression of TWIST1 by CRISPR/Cas9 results in a reduction of tumor growth and metastasis colonization in immunocompromised mice. Moreover, TWIST1 knockout tumors display a less aggressive cellular morphology and a reduced disruption of the extracellular matrix (ECM) reticulin network. Additionally, we identify a TWIST1-mediated transcriptional program associated with dismal outcome in NB and involved in the control of pathways mainly linked to the signaling, migration, adhesion, the organization of the ECM, and the tumor cells versus tumor stroma crosstalk. Taken together, our findings suggest TWIST1 as novel therapeutic target in NB.

## Introduction

Neuroblastoma (NB) is the most prevalent solid extra cranial tumor of childhood [1]. While it accounts for approximately 5% of all pediatric cancer, it contributes for 12% of all pediatric deaths [2, 3]. Primary tumors can arise along the sympathetic chains and in the adrenal medulla [1, 4]. NB is both biologically and clinically heterogeneous. It spans from tumors with favorable biology that can spontaneously regress, to high-risk (HR) disease frequently relapsing or refractory to multimodal treatments and responsible for 50-60% of mortality [1, 4]. Prognosis is associated with a number of factors, including International Neuroblastoma Risk Group (INRG) stages, age at diagnosis, histopathological classification, the presence of segmental chromosomal alterations [1, 5], the activation of telomere maintenance mechanisms [6, 7] and somatic mutations in the RAS/MAPK and p53 pathway [7].

Amplification of MYCN (MNA), present in 20% of primary NB and in 40-50% of HR cases, still remains the most important biological predictor of a poor outcome [2].

As for most pediatric cancers, the origins of NB can be linked back to defects in key cell signaling pathways during embryonic development [8]. NB originates from trunk neural crest (NC) progenitors committed to give rise to the sympathetic nervous system [4, 8]. NC cells are a transient population of multipotent cells that, in the developing embryo upon an epithelial to mesenchymal transition (EMT), delaminate, migrate and differentiate into a broad lineage repertoire [9].

TWIST1/2 transcription factors are among the master regulators of the EMT process [10, 11]. TWIST1/2 are highly conserved and guide developmental programs including cell lineage determination and differentiation, and are essential for organogenesis [10, 12]. Reactivation and aberrant functions of TWIST1/2 have been found in several carcinomas. Both TFs provide cells with critical properties including self-renewal capabilities, resistance to oncogene-induced failsafe programs and invasive capabilities thus promoting cancer initiation and progression toward a metastatic disease [10, 11, 13]. Since TWIST1/2 are active in NC cells, where they play a key role in driving EMT and migration, the study of their functions in NB is particularly important to better understand the neuroblastomagenesis, as distant metastases are already present by the time of diagnosis for the disseminated forms of this disease. So far, the role of TWIST1/2 in NB is still largely unknown. Upregulation of TWIST1 is found in NB with MNA and in a subset of no-MNA tumors, overexpressing MYCN or MYC [14–16]. In addition, TWIST1 protects NB cells from the pro-apoptotic effects mediated by MYCN, through the inhibition of the ARF/p53 pathway and cooperates with MYCN in NB to uphold both *in vitro* cell proliferation and *in vivo* tumor growth [14, 17]. Recently, TWIST1 was also identified as a key regulator of MYCN-driven gene regulation through their cooperative binding on enhancers [17].

In this study, we initially revealed the correlation between the expression of TWIST1 and NB clinical prognostic factors *in silico* on primary NB gene expression datasets and in tumor tissue microarrays. Using an *in vivo* model for transcriptomic analyses, we then unveiled the impact of CRISPR/Cas9-mediated TWIST1 silencing on NB tumor growth, metastatic colonization and the reorganization of the tumor microenvironment (TME).

## Methods

### Tumor Microarray (TMA) and Immunohistochemistry

The TMA was composed by 72 primary tumors, 25 matched metastases and 44 matched control normal tissues (13 sympathetic ganglia and 31 adrenal glands, Supplementary Table S1) obtained from 72 patients diagnosed with NB between July 1988 and November 2001 treated and followed at the Bicêtre hospital (Le Kremlin-Bicêtre) and the Gustave Roussy Institute (Villejuif).

Immunohistochemical study on patient tissues was performed after patients’ informed consent and according to the ethical regulations of the institution. On average, 4 tissue cores with a 0.6 mm diameter were obtained and transferred into a recipient paraffin block using a tissue arrayer (Alphelys: Beecher Instruments Micro-Array Technology, Plaisir, France). TMA sections 5-μm were made on Benchmark XT Ventana (ROCHE Diagnostics). After dewaxing, antigen retrieved is performed using water-bath heating in the following buffers: in citrate buffer pH 6.0 (CC2 citrate-based buffer Ventana Medical Systems ROCHE Diagnostics) for TWIST1 and in a CC1 buffer of pH 8 (CC1 = Tris-Borate/ EDTA, Ventana Medical Systems ROCHE Diagnostics) for TWIST2. Slides were then incubated 1h at RT with the rabbit polyclonal antiTWIST1 (1/50, ABD29, Millipore, Burlington; MA, USA); or 1h at 37°C with the sheep polyclonal anti-TWIST2 (1/200, AF6249, R&D Systems, Minneapolis, MN, USA) in Antibody Diluent Buffer from Ventana Medical Systems, ROCHE Diagnostics. The detection kit for the antibodies is the UltraView DAB detection Kit (Ventana Medical Systems Inc./ Roche Diagnostic). A counter-staining of the nuclei was used for 12 minutes by Hematoxylin. Immunostaining scores (0–4) were established for each stained tissue by semi-quantitative optical analysis by two independent investigators blinded for clinical data. The percentage of positive cells in each sample was scored as follows: 0, all cells negative; 1+, up to 25% of cells were positive; 2+, 26% to 50%; 3+, 51% to 75%; 4+, more than 75%.

### Cell culture

The established human MNA NB cell lines (SK-N-Be2c and LAN-1) were obtained from their lab of origine [18, 19]. Authentication of SK-N-Be2c and LAN1 cell lines was performed by microsatellite short tandem repeat analysis before starting the transduction experiments (Microsynth, Switzerland). The no-MNA NB1-M primary cells were derived in our laboratory from a bone marrow tissue recovered at the diagnosis from a patient with NB at the Hematology Oncology Unit of the University Hospital of Lausanne, Switzerland [20]. All cell lines were cultured in Dulbecco’s modified Eagle’s medium (D-MEM) (Gibco, Paisley, UK), supplemented with 1% penicillin/streptomycin (Gibco) and 10% heat inactivated Fetal Calf Serum (FCS) (Sigma-Aldrich, St. Louis, Missouri, USA) and under standard culture conditions in humidified incubator at 37°C with 5% CO_2_.

### In vivo studies

Animal experiments were carried out with athymic Swiss nude mice (Crl:NU(Ico)-Foxn1^nu^; Charles River Laboratory, France) in accordance with established guidelines for animal care of the Swiss Animal Protection Ordinance and the Animal Experimentation Ordinance of the Swiss Federal Veterinary Office (FVO). Animal experimentation protocols were approved by the Swiss FVO (authorization numbers: VD2995 and VD3372). All reasonable efforts were made to reduce suffering, including anesthesia for painful procedures. For surgical procedures, mice were anaesthetized using isoflurane (Baxter, Deerfield, IL, USA) and received paracetamol as analgesia the day before the surgery. Orthotopic implantations were performed as previously described [21] with slight modifications: 5×10^5^ (ortho_1, 6 mice/group) and 5×10^4^ (ortho_2, 12 mice/group) SK-N-Be2c cells were resuspended in 10 μl of PBS and injected in the left adrenal gland after a small incision above the left kidney. Tumor growth was followed by ultrasound every 7 to 14 days at the Cardiovascular Assessment Facility (University of Lausanne). For subcutaneous implantation, groups of 5 mice were injected in the right flank with 2.5×10^5^ cells suspended in 200 μl 1:1 mix of DMEM and BD Matrigel^™^ Basement Membrane Matrix (BD Biosciences, Bedford, MA, USA). The grafted animals were then weekly monitored with calipers for tumor growth assessment. The tumor volume was calculated using the formula: volume = 4/3 × π × (depth × sagittal × transversal)/6 for ortho tumors; and volume = (length x width^2^)/2 for sc tumors. For both orthotopic and subcutaneous implantations, mice with tumor volumes around ~1000 mm^3^ were sacrificed using CO_2_. Tumors and organs (lungs, liver) were cut into pieces and snap frozen in liquid nitrogen or fixed in formol and embedded in paraffin (lungs, liver, kidneys and spleen).

### RNA isolation

Total RNA from cell lines and tumors was extracted using RNeasy kit (Qiagen, Hilden, Germany). RNA concentration was quantified using a Nanodrop (Agilent Technologies, Wilmington, DE, USA). For the RNA sequencing, RNA was quantified using Qubit Fluorometer (Life Technologies, Carlsbad, CA, USA).

### RNAseq library preparation

RNAseq was performed at the iGE3 Genomics platform (University of Geneva, https://ige3.genomics.unige/ch) using standard techniques RNA integrity was verified using the Agilent 2100 Bioanalyzer system (Agilent Technologies). The total RNA ribo-zero gold kit from Illumina was used for the library preparation with 1 μg or 500 ng of total RNA as input for cells (n=3 biological replicates/group) and tumors (n=4/group), respectively. Library molarity and quality were assessed with the Qubit and Tapestation using a DNA High sensitivity chip (Agilent Technologies). Libraries were pooled at 2 nM and loaded for clustering on 1.5 lanes for cells and 1.5 lanes for tumors of a Single-read Illumina Flow cell. Reads of 100 bases were generated using the TruSeq SBS chemistry on an Illumina HiSeq 4000 sequencer.

### Bioinformatics analysis of RNAseq data

For all samples, fastq files with 100 nucleotides long single-end reads were mapped with STAR version 2.5.2b on both the Human genome version Hg19 and the Mouse genome version Mm10, simultaneously. The following options were changed from the default parameters:-outSAMmultNmax 50; --outFilterMatchNminOverLread 0.4; --quantMode TranscriptomeSAM.

Transcriptome annotations in gtf format for both organisms were downloaded from the gencode website (https://www.gencodegenes.org/). Reads mapped on either the Human or the Mouse transcriptome were then parsed and split in one file per organism with an in-house perl script. Reads with matches on both Human and Mouse were discarded from the Mouse file. Per-gene counts and rpkm were then extracted independently for each organism using rsem version 1.3.0. All RNAseq per-gene data quality checks and analysis were done in R. Mouse and Human data were analyzed independently, but following the same protocol. Protein coding genes with a log2(rpkm) value above 1 in at least one sample were kept (13742 genes in SK-N-Be2c for Human data; 14538 for Mouse data). Principal Component Analysis were done using the normalized log2 (rpkm) values. Clustering analysis were performed on the normalized log2 (rpkm) values using euclidean distance measures and the ward.D2 agglomeration method. Differential analyses were performed using the raw counts in DESeq2 package version 1.26.0. For each comparison, the cutoffs for fold-change (in log2) and adjusted *p* values to call differentially transcribed genes were set to 1 and 0.05 for Human, respectively, and to 0.5 and 0.05 for Mouse, respectively. Heat maps for sample correlations and for specific gene lists were generated using the heatmap.2 function from the gplots package version 3.0.1.2 on the log2 of DESeq2 normalized counts. Functional gene ontology analysis was performed by applying a hypergeometric test on selected genes lists against gene sets from KEGG, GO (Molecular Function, Biological Process and Cellular Component), REACTOME, and BIOCARTA pathways. The *p* value cutoff for terms selection was set to 0.001 for Human data and to 0.01 for Mouse data; only those terms with an adj *p* value below 0.01 and 0.1 were taken into consideration for the graphical representation, respectively. For the GO analysis of the secretome, the lines containing multiple gene references were split before to apply the hypergeometric test on the resulting list of genes (673 terms in the secretome vs 678 terms in the transcriptome). For external RNAseq data analysis (Super series number: GSE80154; SubSeries number: GSE80153), fastq files from GSM2572350 to GSM2572355 corresponding to Be2C samples at 0 (DMSO: GSM2572350 to GSM2572352) and JQ1 24h (GSM2572353 to GSM2572355) were downloaded. These samples were then re-analyzed by applying the same protocol used for the local RNAseq data.

### Protein extraction for cell secretome analysis

Three independent conditioned media (CM) samples were recovered from SK-N-Be2c Control and sgTWIST1 cells. Once cells reached ~75% of confluence, the medium was replaced with FBS- and phenol red-free DMEM (Gibco) in which cells were incubated for 24 hours. CM were first clarified by three centrifugation steps: 10’ at 300 x g; 10’ at 2000 x g cells; and 30’ at 10000 x g at 4°C, and then concentrated using 15 ml Amicon spin filter cartridges (cutoff: 3 kDa, 10705884-Merck Millipore, Burlington, MA, USA) by serial addition of 10 ml of CM and centrifugation at 4000 x g until 1.5 ml were left. After dilution in 100 mM Ammonium Bicarbonate buffer to the starting volume, the CM were re-concentrated by centrifugation at 4000 x g, and these steps were repeated twice until 0.5 ml were left. Finally, aliquots were snap frozen in liquid nitrogen and used for the LC-MS analysis performed at the Protein Analysis Facility (University of Lausanne, Switzerland). CM were dried in a SpeedVac and then digested according to a modified version of the iST protocol (61). Pellets were resuspended in 50 μl of modified iST buffer (2% sodium deoxycholate, 20mM DTT, 5mM EDTA, 200mM Tris pH 8.6) and heated at 95°C for 5 minutes. 50 μl of 160 mM chloroacetamide (in 10 mM Tris pH 8.6) were then added and cysteines were alkylated for 45 minutes at 25°C in the dark. After 1:1 dilution with H2O, samples were adjusted to 3 mM EDTA and digested with 0.5 μg Trypsin/LysC mix (Promega #V5073) for 1h at 37°C, followed by a second 1h digestion with a second, identical aliquot of proteases. To remove sodium deoxycholate, two sample volumes of isopropanol containing 1% trifluoroacetic acid (TFA) were added to the digests, and the samples were directly desalted on a strong cation exchange (SCX) plate (Oasis MCX; Waters Corp., Milford, MA) by centrifugation. After washing with isopropanol/1% TFA, peptides were eluted in 250ul of 80% MeCN, 19% water, 1% (v/v) ammonia.

### Mass spectrometry analyses

Tryptic peptides fractions were dried and resuspended in 0.05% TFA, 2% (v/v) acetonitrile, for mass spectrometry analyses. Tryptic peptide mixtures were injected on an Ultimate RSLC 3000 nanoHPLC system (Dionex, Sunnyvale, CA, USA) interfaced to an Orbitrap Fusion Tribrid mass spectrometer (Thermo Scientific, Bremen, Germany). Peptides were loaded onto a trapping microcolumn Acclaim PepMap100 C18 (20 mm x 100 μm ID, 5 μm, 100Å, Thermo Scientific) before separation on a reversed-phase custom packed nanocolumn (75 μm ID × 40 cm, 1.8 μm particles, Reprosil Pur, Dr. Maisch). A flowrate of 0.25 μl/min was used with a gradient from 4 to 76% acetonitrile in 0.1% formic acid (total time: 65 min). Full survey scans were performed at a 120’000 resolution, and a top speed precursor selection strategy was applied to maximize acquisition of peptide tandem MS spectra with a maximum cycle time of 0.6s. HCD fragmentation mode was used at a normalized collision energy of 32%, with a precursor isolation window of 1.6 m/z, and MS/MS spectra were acquired in the ion trap. Peptides selected for MS/MS were excluded from further fragmentation during 60s.

### Mass spectrometry data analysis and processing

Tandem MS data were processed by the MaxQuant software (version 1.6.3.4))[22] incorporating the Andromeda search engine [23]. The UniProt human reference proteome database of January 2019 was used (73’950 sequences), supplemented with sequences of common contaminants. Trypsin (cleavage at K,R) was used as the enzyme definition, allowing 2 missed cleavages. Carbamidomethylation of cysteine was specified as a fixed modification. N-terminal acetylation of protein and oxidation of methionine were specified as variable modifications. All identifications were filtered at 2% FDR at both the peptide and protein levels with default MaxQuant parameters. After inspection and data QC based on the Ibaq [24] values, the LFQ label-free values [25] were used for protein quantitation. MaxQuant data were further processed with Perseus software (66) for the filtering, log2-transformation, normalization of values and the statistical analyses. After removal of contaminants and reverse hits, intensity values were log2 transformed. Only proteins identified by at least two peptides and quantitated in at least all three samples of one condition were retained for further analysis. Missing values were imputed with standard Perseus parameters (normal distribution with width 0.3 and down-shifted by 1.8 SD). An unpaired T-test was used to determine significant changes, corrected for FDR with the Benjamini-Hochberg method and a threshold q-value at 0.01. Imputed values were subsequently removed from tables. Gene Ontology functional analysis were performed as previously described in the “Bioinformatics analysis” section, after splitting the lines containing multiple genes references.

### Statistical analysis

All statistical analyses were performed using GraphPadPrism 8.3.0 (GraphPad Software Inc., San Diego, CA, USA). D’Agostino-Pearson normality test was performed for each data set, and depending on data distribution, they were analyzed with unpaired two-tailed parametric ttest or non parametric Mann-Whitney test to compare two different conditions.

## Results

### High levels of TWIST1 RNA expression are associated with poor outcomes in patients with NB

*In silico* analysis using the CCLE database (https://portals.broadinstitute.org/ccle) shows that NB displays the highest levels of TWIST1 expression among 40 cancer cell lines, whereas TWIST2 is barely detected (Supplementary Fig. S1A). To evaluate whether TWIST1/2 expression correlates with patient outcomes and NB prognostic factors, we analyzed two large clinical cohorts of primary NB tumors using the R2: Genomics Analysis and Visualization Platform (http://r2.amc.nl) (SEQC (19), n = 498; Kocak (20), n = 649). In both datasets, a high level of TWIST1 transcript strongly correlates with both a reduced overall survival (OS) (**Fig. 1A**; Supplementary Fig. S1B) and event-free survival (EFS) (Supplementary Fig. S1C). Moreover, the expression of TWIST1 was more elevated in presence of disease progression (**Fig. 1A**); in MNA NBs (Supplementary Fig. S1D); and in higher stage tumors (stages 3 and 4 vs 1 and 2; stage 4 vs 4s) (Supplementary Fig. S1E). We stratified patients of the SEQC dataset according to the level of TWIST1 expression and either the risk (HR vs low-risk (LR); (**Fig. 1A**) or MYCN status, Supplementary Fig. S1F). For HR or MNA patients, TWIST1 expression level had no impact on the EFS. Conversely, for LR cases and no-MNA tumors, a high level of TWIST1 expression was associated with a reduced outcome, likewise MNA or the HR status, hinting to a possible role for TWIST1 as a prognostic factor of adverse event for these patients. As opposed to TWIST1, in the two same datasets, higher levels of TWIST2 were associated with both a better OS and EFS in NB patients (Supplementary Fig. S1G). Moreover, TWIST2 expression was increased in no-MNA NB (Supplementary Fig. S1H).

**Figure 1.**
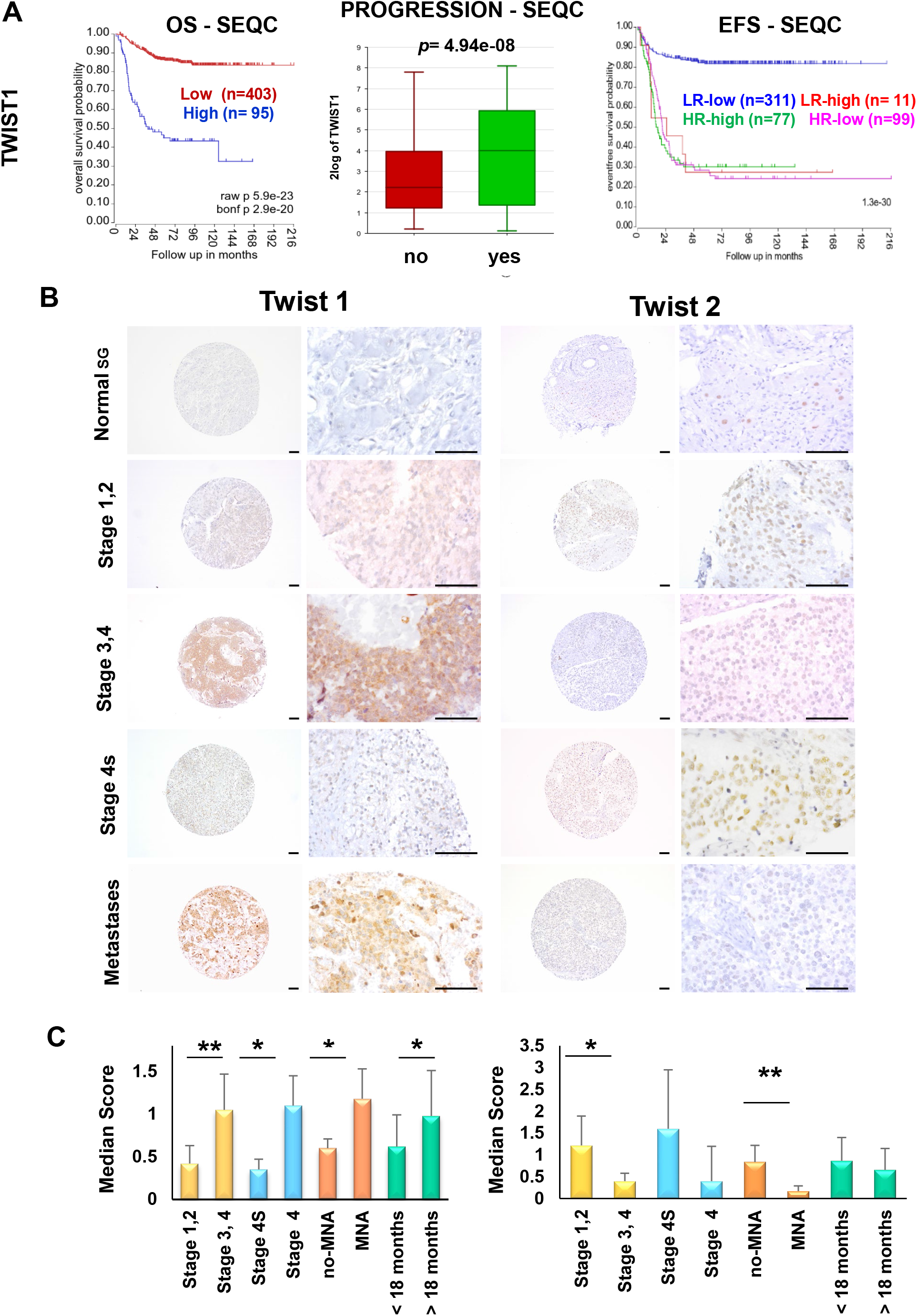
TWIST1 RNA expression is associated with poorer outcome of NB patients and displays an opposite protein expression profile in a NB tissue microarray. (**A**) Analysis of TWIST1 expression in the SEQC dataset of primary NB tumors. Left panel: KaplanMeier OS curve associated with TWIST1 expression. Expression cutoff: 44.441. Middle panel: Box-and whisker plots showing the expression of TWIST1 in relation to disease progression. Right panel: Kaplan-Meier EFS curves showing the stratification of patients of the SEQC dataset according to the risk classification (high-risk: HR; low-risk: LR) and TWIST1 expression (high or low). (**B**) TWIST1 and TWIST2 protein expression were analyzed by IHC using a NB TMA containing 97 tumor sections: 72 primary tumors, 25 matched metastases and 44 matched control normal tissues (i.e. SG). Representative images of TWIST1 and TWIST2 IHC staining are shown for each indicated category. Magnification 100x (left panels) and 400x (right panels); scale bares=100 μm. (**C**) Bar graphs showing the median scores (ms) ± SD of TWIST1and TWIST2 IHC staining for different comparisons (see Table 1). Statistical analysis was done using parametric Student’s t-test.

### TWIST1 expression patterns reveal a correlation with poor prognostic factors in NB

We examined the expression levels of TWIST1/2 proteins in a NB tissue microarray (TMA) (Supplementary Table S1). In control sympathetic ganglia (SG), TWIST1 was not detected while TWIST2 was present with moderate intensity in 46% of SG (**Fig. 1B**; Supplementary Table S1). TWIST1 expression was statistically significantly higher in tumors associated with poor prognosis: stages 3-4 vs stages 1-2; stage 4 vs 4s; tumors with MNA vs no-MNA; and in patients older than 18 months at the diagnosis (**Fig. 1C**; Supplementary Table S1). On the other hand, the expression of TWIST2 was higher in tumors with better prognosis: stages 1-2 vs stages 3-4 and in patients with no-MNA vs MNA (**Fig. 1C**; Supplementary Table S1). However, no statistically significantly differences in TWIST2 expression were observed in stage 4s vs stage 4 or in relation with age at diagnosis (**Fig. 1C**). Finally, TWIST1 was frequently expressed in metastases (76% positive, median score=0.95), while TWIST2 expression was uncommon (30% positive, median score=0.31) (**Fig. 1B**; Supplementary Table S1).

### TWIST1 KO impairs the neurosphere-forming ability of NB cells

To investigate the contribution of TWIST1 in the aggressive features of NB, three cell lines, either MNA (LAN-1 and SK-N-Be2c) or non-NMA (NB-1), were chosen for a TWIST1 knockout (KO) through CRISPR/Cas9. A complete KO of the wild type (wt) TWIST1 protein expression was obtained with the sgTWIST1 #1 for the three cell lines that from now on will be referred to as sgTWIST1 cells (Supplementary Fig. S2A and B). TWIST1 KO did not significantly affected the 2D growth property of NB cell lines (Supplementary Fig. S2C), however it reduced the neurosphere-forming ability of the three NB cell lines (**Fig. 2A**). Consequently, the number of sgTWIST1 cells recovered from primary neurospheres was statistically significantly lower compared to Control cells (**Fig. 2A**), indicating the role played by TWIST1 in propagating a highly tumorigenic subpopulation of NB cells.

**Figure 2.**
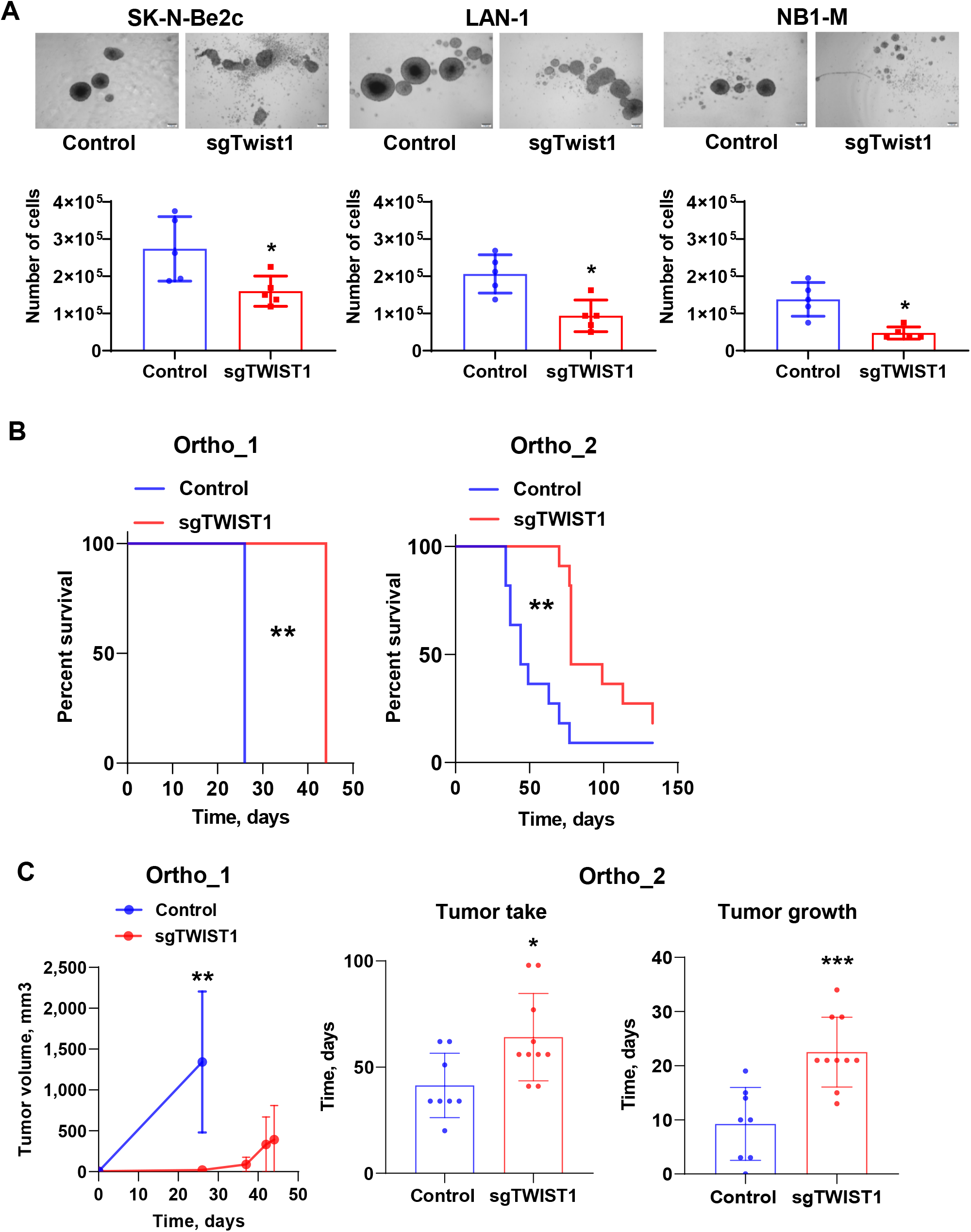
TWIST1 KO reduces the neurosphere forming capacities of NB cells *in vitro* and the tumor growth capacities of SK-N-Be2c cells *in vivo*. (**A**) Upper panel: representative images (scale bar 200 μm) showing the size and shape of primary neurospheres of Control and sgTWIST1 NB cells after 7 days in culture. Lower panel: the numbers of cells obtained after dissociation of Control and sgTWIST1 primary neurospheres are plotted in bare graphs as individual values for each independent experiments and mean ± SD (n=5 experiments performed in duplicates). Mann Whitney test: \**p*=0.0317 for SK-N-Be2c; \**p*=0.0159 for LAN-1 and NB1-M. (**B**) Kaplan-Meier survival curves of athymic Swiss nude mice implanted orthotopically with SK-N-Be2C-Control or -sgTWIST1 cells. Mice were sacrificed once tumors reached the volume of 1000 mm^3^ and 500 mm^3^ for ortho_1 and ortho_2 experiments, respectively. Tumor take: ortho_1: 100% (6/6) in the Control group, 66.66% (4/6) in the sgTWIST1 group; ortho_2: 89% (8/9) in the Control group, 83% (10/12) in the sgTWIST1 group. Median survival in the Control vs sgTWIST1 group: 26 vs 44 days for ortho_1 (\*\**p*=0.0027); 49 vs 78 days for ortho_2 (\*\**p*=0.0016). Gehan-Breslow-Wilcoxon test. (**C**) Left panel: Tumor growth (mean tumor volumes ± SD) for ortho_1 experiment. Multiple t-test (HolmSidak, α=0.05, without assuming a consistent SD): \*\**p*=0.0037. (Middle and right panel: Time for tumor initiation (middle) and tumor growth (right) in the ortho_2 experiment (mean days ± SD). Tumor initiation correspond to the number of days required to measure an AG volume > 10 mm^3^ (mean Control: 41.38 days, sgTWIST1: 64.10 days, \**p*=0.0192). Time for tumor growth was calculated as the number of days at sacrifice minus the number of days for tumor initiation (mean Control: 9.25 days, sgTWIST1: 22.50 days, \*\*\**p*=0.0006, unpaired t-test).

### TWIST1 KO delays tumor growth of NB xenotransplantation and extends survival in mice

Next, we investigated the contribution of TWIST1 in the tumorigenicity of NB cells. In three independent experiments, athymic Swiss nude mice were injected with the SK-N-Be2C Control and sgTWIST1 cells either orthotopically (500’000 cells for ortho_1 and 50’000 cells for ortho_2) or subcutaneously (sc, 250’000 cells). In all the three models, the growth of the sgTWIST1 tumors was severely delayed compared to Controls thus extending sgTWIST1 mice survival (**Fig. 2B**, Supplementary Fig. S3A). In particular, in the first orthotopic experiment (ortho_1), 26 days after the injection, tumors in Control mice were already above the predetermined volume for sacrifice while the sgTWIST1 mice were still in the lag phase (**Fig. 2C**). In the second orthotopic experiment (ortho_2), we observed a significant delay for both SK-N-Be2c-sgTWIST1 tumor initiation and tumor growth (**Fig. 2C**). Furthermore, 25 days after sc injections, the size of Control tumors was ~10 times larger than sgTWIST1 tumors, which required four additional weeks to grow (Supplementary Fig. S3B). Finally, in both orthotopic experiments we observed SKN-Be2c-Control tumors invading the vena cava (n=3/6: ortho_1; n=3/8: ortho_2) (Supplementary Fig. S3C), whereas no invasion was detected in the sgTWIST1 mice group.

### TWIST1 KO diminishes the malignant phenotype of tumors and decreases intrapulmonary macrometastasis

In both orthotopic *in vivo* models, Control tumors presented histological features corresponding to undifferentiated or poorly differentiated cells, while sgTWIST1 tumors were more differentiated (Fig. 3A, left panel). Moreover, Control cells showed a lower degree of cohesion and a higher degree of immune cell infiltration compared to the sgTWIST1 tumors (Fig. 3A, left panel). We analyzed the effects of TWIST1 KO on the pattern of collagen III/reticulin fibers, which contribute to the ECM. Throughout all the three *in vivo* models, in Control tumor tissues the continuity of the reticular fiber framework was lost in extensive tumor areas, and we observed irregular thickening and fraying of fibers mainly at the borders of tumors (**Fig. 3A**, Supplementary Fig. S3D). In contrast, the sgTWIST1 tumors were characterized by a preserved reticulin mesh, resembling that of the normal adrenal gland (AG) (**Fig. 3B**, Supplementary Fig. S3D). This effect was not altered by tumor size at sacrifice (Supplementary Fig. S3E).

**Figure 3.**
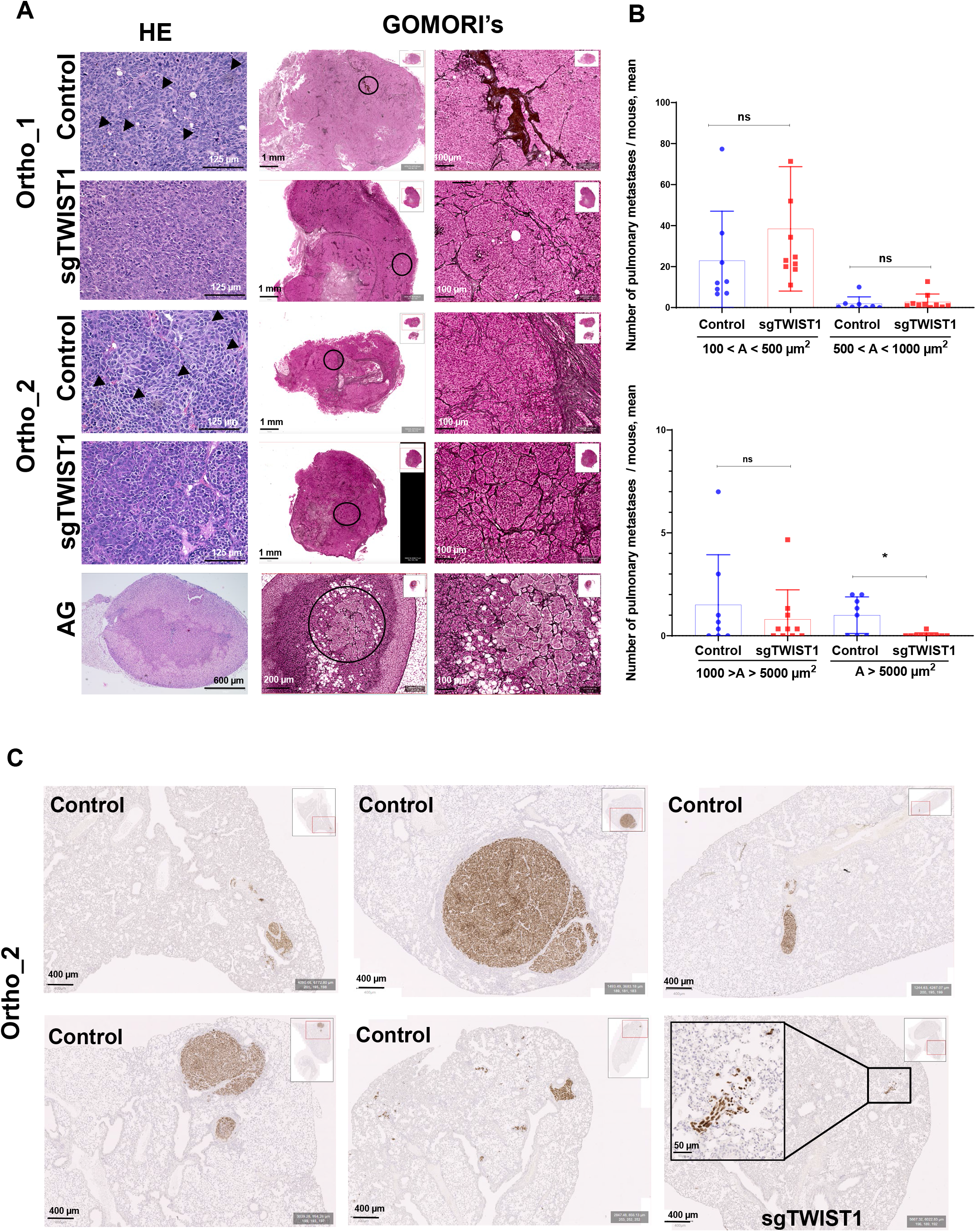
TWIST1 KO produces tumor with a less aggressive phenotype and impairs the formation of the intrapulmonary macrometastases. (**A**) Left panel: representative images of H&E staining of ortho tumors and AG. H&E staining of both ortho-derived tumors depicted cells in control tissues separated by thin fibro-vascular septs having irregular size and shape; no discernable/scarce cytoplasm; one or few prominent nucleoli; spindle-shaped cells with fusiform nuclei (black arrow) that tended to have a fascicular organization. Conversely, sgTWIST1 tumor cells were portrayed by a more regular size and shape (round to oval) with only slight irregularities, finely granular (“salt-and-pepper”) chromatin, small nucleoli and moderate/more discernible cytoplasm (scale bar: 125 μm for tumors; 600μm for AG). Middle and right panels: representative images of Gomori’s staining showing the architecture of the collagen III/reticulin fibers in ortho tumors and AG. Middle panels: large views of tumor and AG sections; scale bars: 1 mm and 200 μm, respectively. Right panels: zoomed view of the region highlighted by a black circle, scale bars: 100 μm for both tumors and AG. **B**) Quantification of metastases detected by IHC with the Alu positive probe II within the parenchyma (intrapulmonary) of mice. Data are plotted in a bar graph showing individual values and mean ± SD for micrometastases (upper panel: 100-500 μm^2^: *p*=0.1120; 500-1000 μm^2^: *p*=0.3705) and for macrometastases (lower panel: 1000-5000 μm^2^, *p*= 0.5724; >5000 μm^2^, \**p*= 0.0178). Mann-Whitney test. Percent of mice with macrometastases = 62.5% in the Control group; 10% in the sgTWIST1 group (*p*=0.043 Fisher’s exact test).(**C**) Representative images of Alu positive probe II staining of lungs of the 5 Control and 1 sgTWIST1 ortho_2 mice with pulmonary metastases A > 105 μm2.

Such ECM modifications associated with TWIST1 expression could be responsible for a “pro-neoplastic” stromal phenotype, offering less resistance for the invasive cells to escape the primary tumor site and form metastasis [26]. Therefore, the lungs of the ortho_2 experiment mice were analyzed for the presence of intrapulmonary metastasis. No differences in the number of intrapulmonary micrometastases (area (A) <1000 μm^2^) and in macrometastases with A< 5000 μm^2^ were observed between the two group of mice (**Fig. 3B**). Conversely, the number of intrapulmonary macrometastases with A> 5000 μm^2^ in the sgTWIST1 mice was statistically significantly reduced as a single one was detected in only 1/10 sgTWIST1 mouse (10.7 x10^3^ μm^2^), whereas 5/8 Control mice had multiple macrometastases (**Fig. 3B, C**).

### Identification of distinct transcriptional program regulated by TWIST1 and MYCN in NB cells

Transcriptomic analyses of SK-N-Be2c-Control and –sgTWIST1 cells and their derived ortho_1 tumors were performed by RNAseq. Principal Component Analysis (PCA) revealed a high degree of segregation of the transcriptomic profiles of Control and sgTWIST1 for both cells and ortho_1 tumors, enabling the accurate identification of genes that are differentially expressed (DE) (**Fig. 4A**). We identified 2342 DE genes (1401 up- and 941 down regulated) in SK-N-Be2c cells and 2013 (1003 up- and 1010 down regulated) in the SK-N-Be2c ortho_1 tumors, with 1213 found in common (**Fig. 4A**; Supplementary Fig. S4A; Supplementary Table S2). Gene ontology (GO) analyses for the DE genes in cells and in tumors reported a number of significantly enriched terms related to signaling, nervous system development, migration, proliferation, ECM organization and adhesion for both biological processes (BP) and cellular components (CC) (**Fig. 4B**; Supplementary Fig. S4B; Supplementary Table S3).

**Figure 4.**
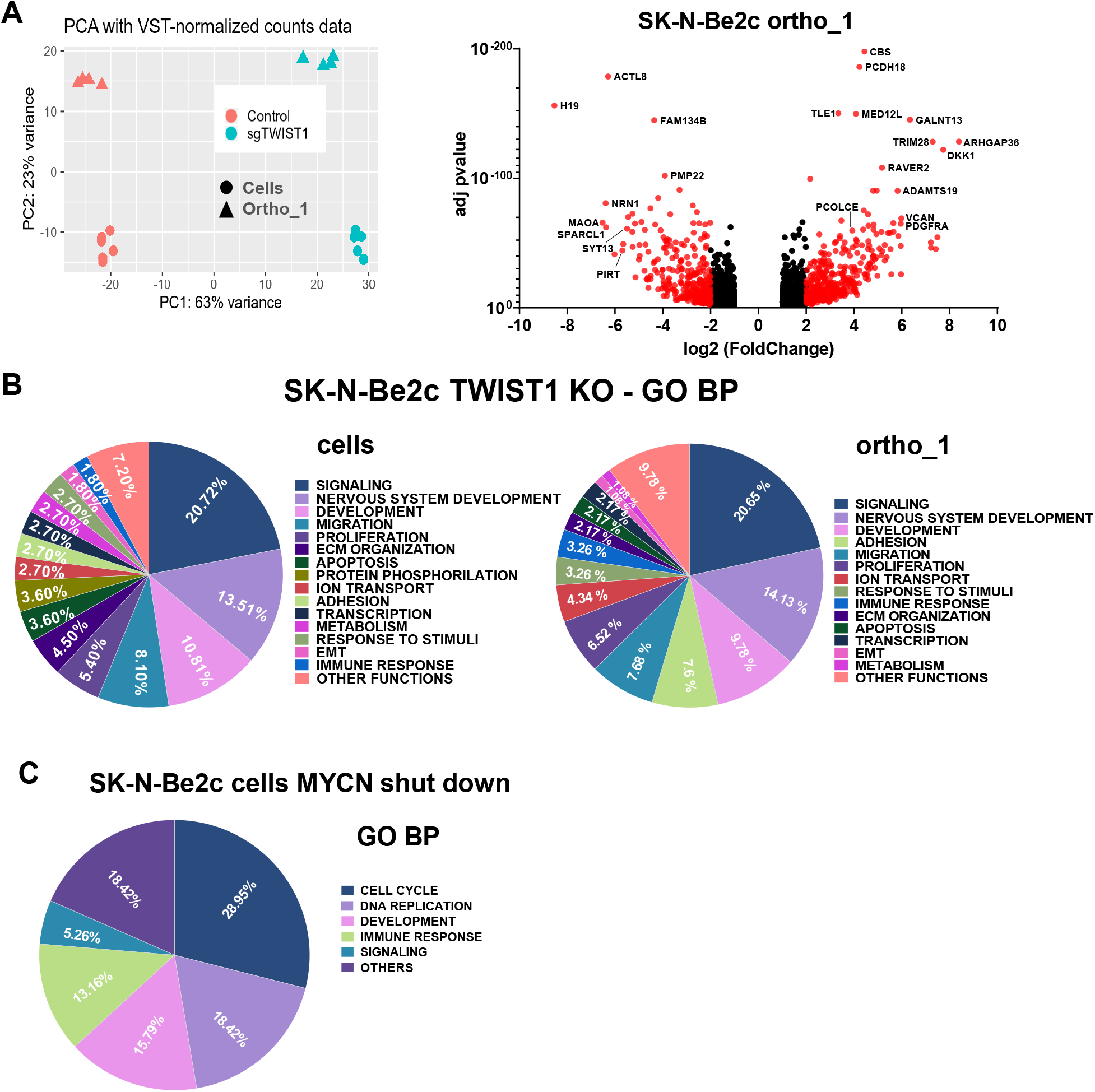
The biological pathways deregulated by TWIST1 KO are distinct from those mediated by MYCN shut down. (**A**) Left panel: PCA samples repartition using the VST-normalized counts. PCA1 and PCA2 are 63% and 23% of total variation, respectively. Right panel: volcano plots showing the distribution of the DE genes according to FC (log2) and adj *p* value between the SK-N-Be2c-Control and –sgTWIST1 ortho_1-derived xenografts. Genes with False Discovery Rate (FDR) < 0.05 and absolute value (av) of log2(FC) ≥ 1 were considered as DE; in red genes with av of log2(FC) ≥2, in black genes with av of log2(FC) ≥ 1 and <2. Positive and negative x-values represent genes either up- or down-regulated by TWIST1, respectively. (**B**). Illustration of the biological processes gene sets found enriched by GO analyses (GO BP) in the DE genes following TWIST1 KO for both SK-N-Be2c cells (left panel) and ortho_1 tumors (right panel). Data are reported as the repartition (in %) of the diverse pathways identified with a FDR < 0.01 (n=111 for cells, n=92 for tumors). (**C**) Illustration of the GO BP gene sets found enriched in the DE genes in SK-N-Be2c cells upon JC1-medited MYCN shutdown. RNAseq data of SK-N-Be2c cells treated with JC1 during 24h or DMSO as control were uploaded (GSE80154, see Methods) (Zeid et al.). Genes with False Discovery Rate (FDR) < 0.05 and absolute value (av) of log2(FC) ≥ 1 were considered as DE. Data are reported as the repartition (in %) of the diverse pathways identified with a FDR < 0.01 (n=38).

As downregulation of MYCN was observed upon transient TWIST1 silencing in SK-N-Be2c, a decrease in MYCN expression level could be, in part, responsible for the deregulation of the transcriptional program observed in our ortho tumors [17]. To exclude this possibility, we analyzed the expression level of MYCN protein by immunoblotting in tumors coming from the three in vivo experiments. In all sgTWIST1 tissues, we detected an increase in the level of MYCN protein compared to the Control counterpart (Supplementary Fig. S4C) although this increase was not sufficient alone to promote and sustain a more aggressive phenotype in the sgTWIST1 tumors.

To compare the transcriptional program defined by TWIST1 with the one induced by MYCN in SK-N-Be2c cells, we reanalyzed RNAseq data obtained upon MYCN shutdown using the BET bromodomain inhibitor JC1 [17]. GO analyses performed on DE genes highlighted an enrichment of gene sets mainly involved in the regulation of cell cycle and the DNA replication for both BP and CC, thus suggesting distinct functions for the two TFs (**Fig. 4C**; Supplementary Fig. S4D; Supplementary Table S4 and S5).

### A TWIST1-mediated gene expression signature is associated with poor outcome in NB

To identify a TWIST1-associated gene signature relevant in primary NB we combined our ortho_1 transcriptomic analysis with RNAseq data of primary NB tumors. Using the ‘R2 Platform, we first listed the genes either correlated (R positive) or anti-correlated (R negative) with TWIST1 expression in the SEQC dataset of NB tumors (n=7737 genes with R absolute value >0.225). Second, we crossed this list of genes with the 2011 DE genes between SK-N-Be2c-Control and -sgTWIST1 tumors, either up-(FC positive) or downregulated (FC negative) by TWIST1. We found 763 genes in common (**Fig. 5A**; Supplementary Table S6) among which we selected those that had both R and FC either positive (172 genes) or negative (317 genes). We called these resulting 489 genes the TWIST1-signature (**Fig. 5A**; Supplementary Table S6). Using the same SEQC dataset, we analyzed the clinical significance of the signature, and observed that genes correlated with TWIST1 in NB patients and upregulated by TWIST1 in ortho_1 tumors (R and FC positive) mostly had an elevated level of expression in high-risk, more advanced stages and MNA tumors. In addition, these tumors displayed a low level of expression of genes downregulated in the TWIST1-signature (**Fig. 5B**). Finally, an elevated expression level of the TWIST1-signature was associated in the SEQC and Kocak datasets with a poor OS and EFS for both the complete patient cohorts and the sub-cohorts without MNA (**Fig. 5C**).

**Figure 5.**
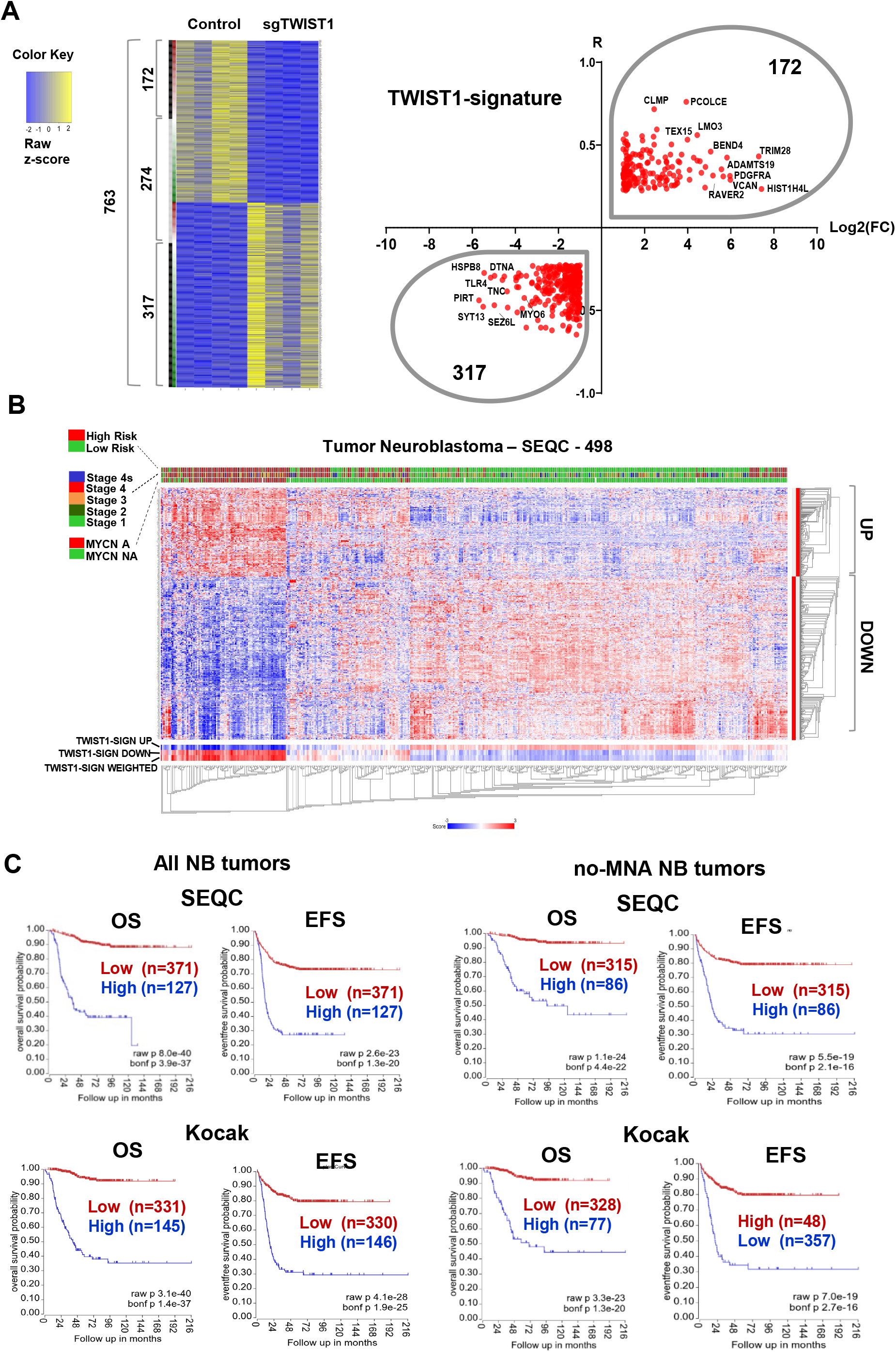
Identification of a TWIST1-associated gene signature correlating with poor prognosis in NB. (**A**) Left panel: heatmap showing 763 common genes either correlated or anticorrelated with TWIST1 in NB patients and DE in the ortho_1 tumors. The binary side color bar going from green to red indicates DE genes anti-correlated (R<-0.225, green) or correlated (R>0.225, red) with TWIST1 in the SEQC dataset; the black bar shows the genes that have both FC and R values either positive or negative representing the TWIST1-signature, and the grey bar the genes that have opposite FC and R values (not included in the signature). Right panel: volcano plot showing the distribution of the 489 genes of the TWIST1-signature according to their log2(FC) in SK-N-Be2c ortho_1 tumors and R values in the SEQC dataset. (**B**) Heatmap hierarchical clustering showing different expression pattern relative to TWIST1-signature genes generated using the R2 Platform (http://r2.amc.nl). Columns represent patients annotated in the SEQC cohort; the 489 genes are clustered hierarchically along the left y-axis. Clinical criteria taken into consideration (risk groups, tumor stages, and MYCN amplification status) are indicated on the top by color codes. The heat map indicates in red, blue and white a high, low and a medium level of gene expression (z-score), respectively. The blue-white-red color bars depicted at the bottom of the heatmap represent the z-score of TWIST1_Up and TWIST1_Down gene sub-lists of the signature, as well as for the z-score of the whole signature (weighted). (**C**) Kaplan-Meier OS and EFS survival curves according to the expression level of the TWIST1-signature in both the SEQC and Kocak datasets. Left panel: complete cohort; right panel: sub-cohorts of patients without MNA (no-MNA). Expression cutoff in the SEQC: 0.20 for OS curves; −0.05 for EFS curves. Expression cutoff in the Kocak: 0.03 for all curves.

Among the top deregulated genes in the TWIST1-signature, several have crucial roles during embryonic development, in particular for the correct development of the nervous system (*BMP7*, *FGF2*, *DTNA*, *MATN2*, *PCDHA1*, *PMP22*, *SCL1A3*). Moreover, most of the top upregulated genes are involved in the organization of both TME (*PDGFRA*, *VCAN*, *BMP7*, *FGF2*) and ECM (*ADAMST19*, *PCOLCE*); in the EMT process (*BMP7*, *TRIM28*), as well as in cell proliferation (*FGF2* and *PDGFRA*) and apoptosis (*BMP7*) (**Fig. 6A**). Besides, among the top genes down-regulated in the TWIST1 signature, some are involved in neuronal differentiation (*PIRT*), and various are tumor suppressor genes (*SYT13*, *FAM134B*, *PMP22*, *C7* and *MATN2*) (**Fig. 6A**). Several transcripts belonging to the TWIST1-signature were chosen, based on their degree of differential expression (**Fig. 6A**) and their biological function, for validation by RT-qPCR and WB/IHC. We confirmed that in our xenografts RNA and/or protein levels for *VCAN*, *PDGFRA*, *TRIM28*, *PCOLCE* and *ADAMTS19* were upregulated by TWIST1 while *PIRT* and *SYT13* were downregulated (**Fig. 6B, C**; Supplementary Fig. S5 and S6).

**Figure 6.**
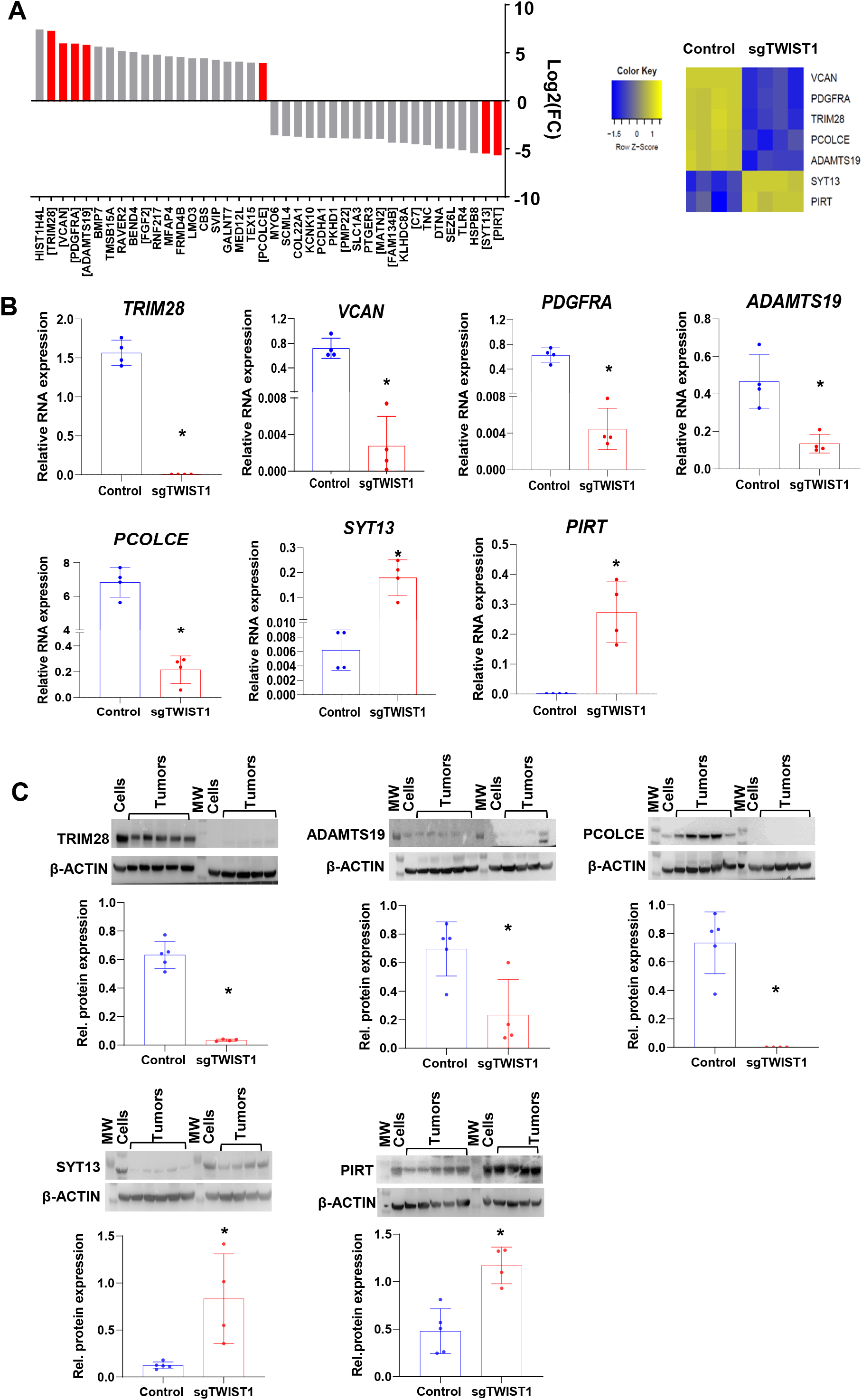
Validation of TWIST1-mediated deregulation of selected genes belonging to the TWIST1 signature in the ortho_1 tumors. (**A**) Left panel: bar plots showing the distribution of the top 20 up- and 20 down-regulated genes of the TWIST1 signature ordered according to their log2(FC). In black, genes that were selected for the validation at both RNA and protein levels. Gene names in brackets indicate up-regulated genes involved in the EMT process, TME organization, proliferation and apoptosis; and down-regulated genes that are known to be tumor suppressor genes or associated with good prognosis in NB. Right panel: heatmap showing the relative RNA expression (z-score) determined by RNAseq of the selected genes in ortho_1 tumors. (**B**) RNA expression levels of the TWIST1 target genes relative to the reference gene *HPRT1* in the ortho_1 tumors analyzed by RT-qPCR are plotted as individual values with mean ± SD. Control n= 6; sgTWIST1 n= 4. Mann Whitney test: \**p*=0.0286 for all comparisons. (**C**) Upper panel: Immunoblotting for TRIM28, ADAMTS19, PCOLCE, ADAMTS19, SYT13 and PIRT (β-ACTIN as the loading control); lower panel: densitometric quantifications of immunoreactive band densities. Expression relative to β-ACTIN were plotted as individual data with mean ± SD. Control n= 5; sgTWIST1 n= 4. Mann Whitney test: *p= 0.0317 for ADAMTS19; *p= 0.0159 for the other proteins.

### TWIST1 alters the level of expression of genes involved in tumor-stroma crosstalk

Cancer cells establish a reciprocal intercellular signaling network and communicate with stromal and immune cells via the production of soluble paracrine factors and their cognate receptors. This complex signaling network shapes the TME to sustain cancer cell proliferation and invasion. To address whether TWIST1 alters the expression of factors involved in cell-cell communication, DE genes annotated as cytokines, chemokines, growth factors, inflammatory mediators and their receptors, as well as integrin and their ligands were extracted from SK-NBe2c tumor transcriptome. This TWIST1-tumor-stroma signature is composed by 77 DE genes, 33 up- and 44 down-regulated (**Fig. 7A**; Supplementary Table S7). Several play a pivotal role in the regulation of focal adhesion (*EGFR*, *ITGA11*, *ITGA6*, *PDGFRB*); cell migration (*COL5A1*, *ITGAV*, *ITGB3*, *PDGFRB*, *TGFB1*); proliferation (*FGF1*, *FIGF*, *IFI16*); angiogenesis (*ACKR3*, *ACVRL1*, *EGFL7*, *FGF1*, *FGFR2*, *FIGF*); and inflammatory and immune responses (*NGFR*, *TNF*, *TNFRSF1A*, *TNFRSF1B*, *TNFRSF4*, *TNFRSF9*, *TNFSF12*, *TNFSF13*, *TNFSF4*). A high level of expression of the TWIST1-tumor-stroma signature was associated with a poor OS and EFS of NB patients in both the SEQC (**Fig. 7A**) and the Kocak datasets (Supplementary Fig. S7A).

**Figure 7.**
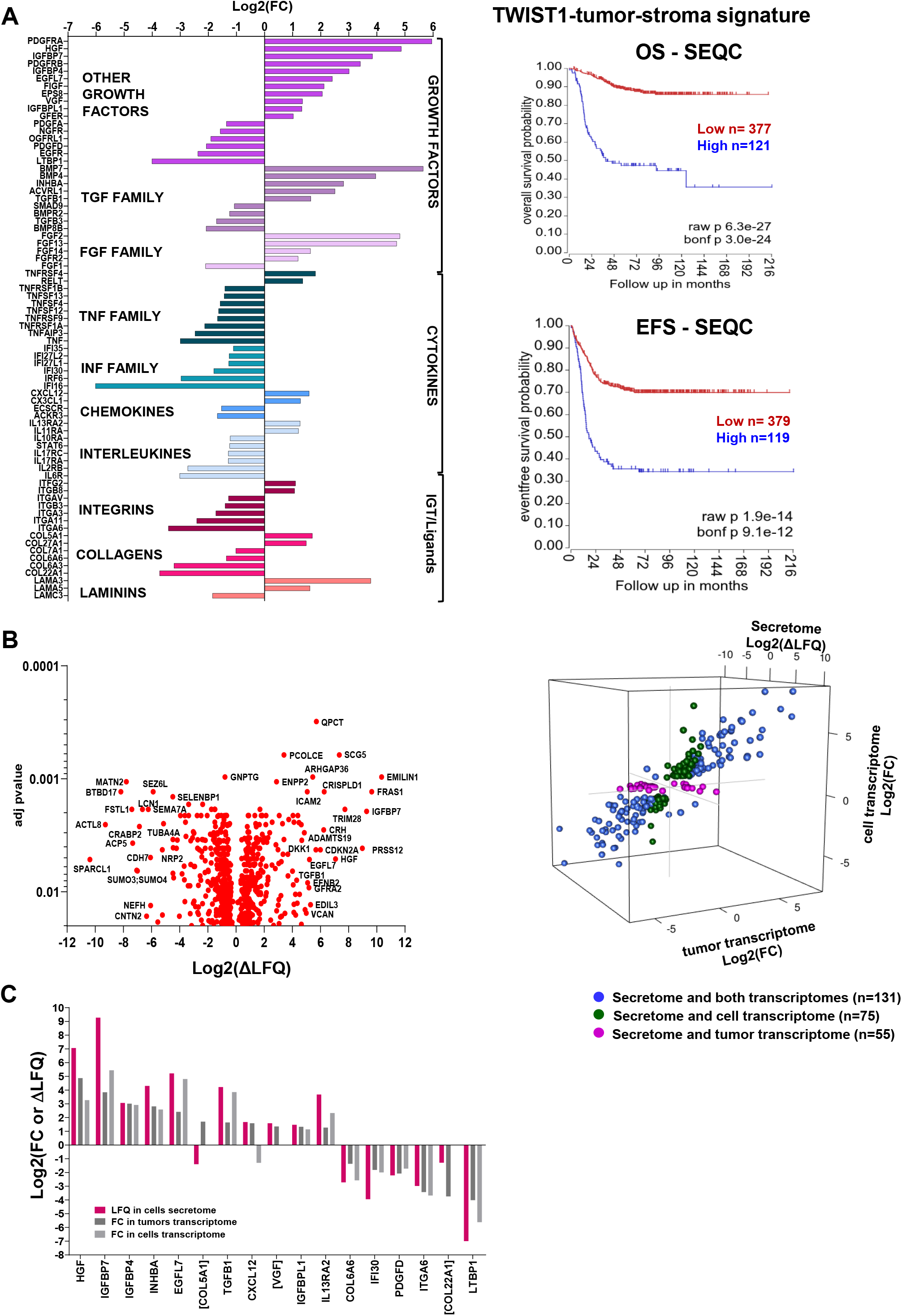
Identification of a TWIST1-mediated-tumor-stroma signature associated with poor outcome in NB. (**A**) Left panel: bar plot illustrating of the 77 DE genes representing the TWIST1-tumor-stroma signature in SK-N-Be2c ortho_1 tumors. Genes were classified according to their log2(FC) in three main categories: growth factors (including the TGF and FGF families cytokines (TNF poor outcome in NB. Right panel: Kaplan-Meier OS and EFS curves of NB patients of the SEQC dataset according to the expression level of the TWIST1-tumor-stroma signature. Expression cutoff for both curves: 0.10. (**B**) Left panel: volcano plot showing the distribution of the DE protein secreted by SK-N-Be2c cells according to the delta label-free quantification (ΔLFQ = LFQ SK-N-Be2c Control – LFQ SK-N-Be2c sgTWIST1) intensities (Log2) and the adjusted p values with an FDR ≤ 0.02 analyzed by LC-MS/MS (n= 3 biological replicates for each group). Right panel: 3D scatterplot showing DE terms in the cell secretome in common with the tumor transcriptome (magenta, n=55), the cell transcriptome (green, n=75), or both transcriptomes (blue, n=131). (**C**) Bar plot showing the terms commonly deregulated between the TWIST1tumor-stroma signature and both the cell transcriptome and secretome. Names in brackets are for terms found to be DE in the secretome but not in the transcriptome of cells.

To validate the tumor-stroma signature at the protein level and further characterize TWIST1-mediated alterations in cell-cell communication, we analyzed the secretome of SK-N-Be2c-Control and -sgTWIST1 cells *in vitro* by HPLC/Tandem MS using their conditioned media (CM) containing both secreted proteins and extracellular vesicles released by tumor cells. These secretomes contained 673 DE peptides (304 up- and 369 down-regulated) (**Fig. 7B**; Supplementary Table S8) that corresponded to 678 proteins. GO analyses revealed an enrichment of BP linked to nervous system development, signaling, response to stimuli, migration, and proliferation (Supplementary Fig. S7B; Supplementary Table S9).

Crossing secretome and transcriptome data from both cells and tumors, we identified 131 commonly deregulated terms, whereas 75 and 55 were uniquely shared between the secretome and either the cell or the tumor transcriptome, respectively (**Fig. 7B**; Supplementary Table S8). Finally, after crossing the TWIST1-tumor-stroma signature with the secretome of cells, we could identify 17 commonly DE terms, among which 14 were also found to be in common with the transcriptome of cells (**Fig. 7C**). Most of the commonly deregulated terms were up regulated by TWIST1 and annotated as growth factors, and for all terms but *COL5A1* and *VGF*, the impact of TWIST1 on RNA and protein expression was always found to be correlated.

### Myofibroblast-associated gene expression is reduced in the stroma of sgTWIST1 orthotopic tumors

Among the terms deregulated in the abovementioned tumor-stroma signature, several are also known for being involved in the crosstalk between cancer cells and the resident and recruited stromal cells (i.e. *TGFB1*, *HGF*, *FGF*, *FGFR*, *EGFR*, *PDGFR*, *CXCL12*) and thus they could mediate a TME sustaining the tumor growth [27]. One of the main stromal changes within a pro-tumorigenic TME is the appearance of cancer-associated fibroblasts (CAFs), playing a critical role in arranging the “soil” within which tumor cells proliferate [28]. To verify whether we could detect the presence of CAFs in the tumor stroma, the ortho_1 RNAseq data were aligned with the murine genome. Between Control and sgTWIST1 tumors, 89 stromal genes were found to be DE (69 up- and 20 down-regulated) (**Fig. 8A**; Supplementary Table S10). Genes up-regulated in the stroma of TWIST1 expressing Control tumors showed a significant enrichment of muscle contraction-related terms (Supplementary Table S11). This was defined as the myofibroblastic signature (n=36 genes) according to the literature [29–32]. GO analysis for the murine DE genes reported a number of statistically significantly enriched terms related to sarcomere organization and muscle contraction (**Fig. 8A**; Supplementary Table S12), supporting a TWIST1-mediated recruitment and activation of myofibroblasts.

**Figure 8.**
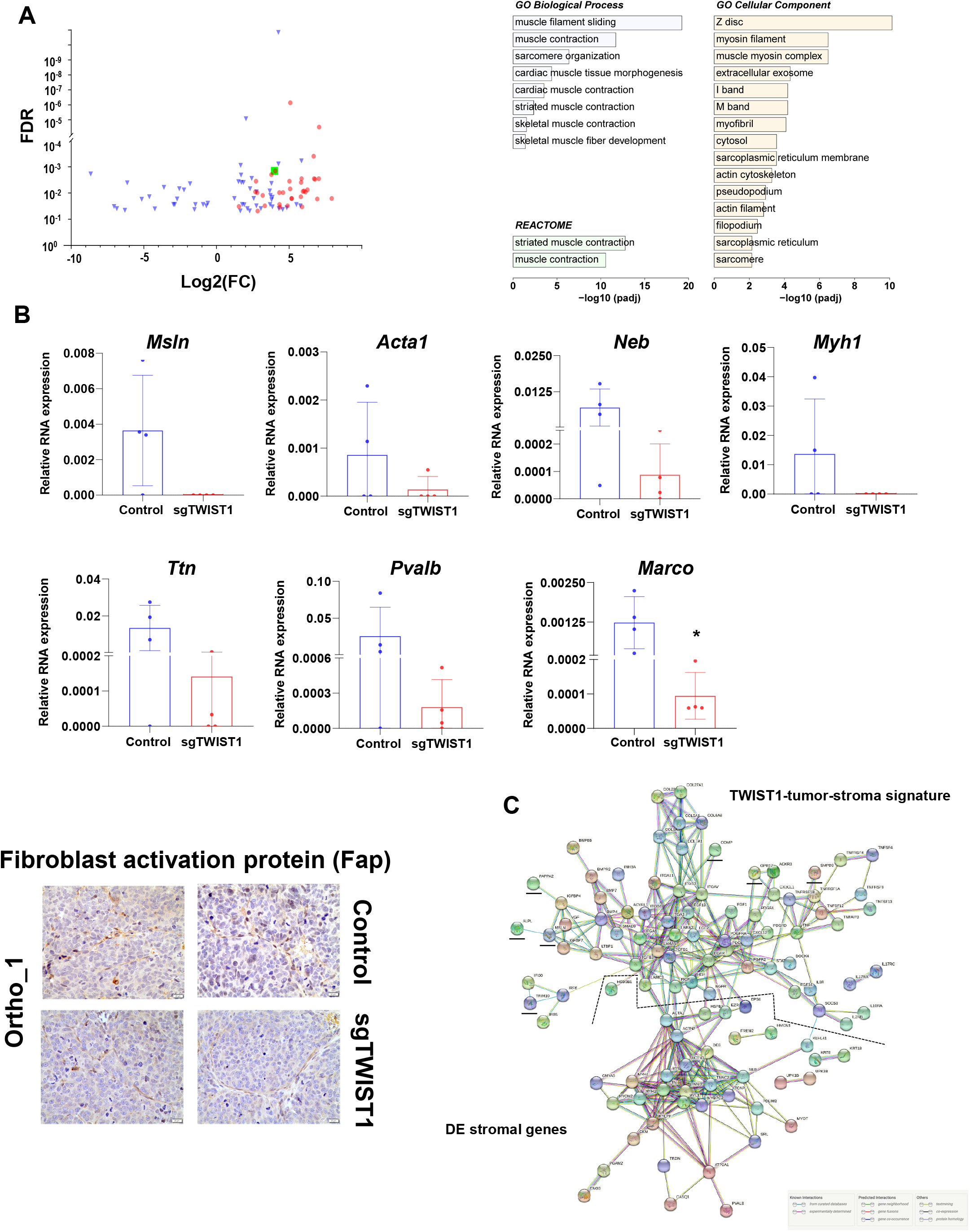
Identification of a TWIST1-associated myofibroblast signature and PPI network for the TWIST1-associated tumor-stroma signature and the DE stromal genes. **(A)** Left panel: volcano plots showing the distribution of the DE gene identified in SK-N-Be2c-Control and –sgTWIST1 tumor stroma of ortho_1 xenografts relative to their log2(FC) and adjusted *p* value (FDR). Genes with FDR < 0.05 and absolute value (av) of log2(FC) ≥ 0.5 were considered as DE. Genes identified as the Myofibroblast signature are indicated in red (n=36). The green square is for the gene *Marco*. Righ panel: bar graph showing the biological processes, cellular components and REACTOME pathways identified by GO analysis of the 89 DE genes of the murine stroma, listed according to their adjusted *p* value. **(B)** Upper panels: mRNA expression levels of the selected myofibroblast genes and Marco relative to β-actin as by RT-qPCR. Data are plotted as individual values with mean ± SD. Mann Whitney test: \**p*= 0.0286. Ortho_1 Control and sgTWIST1 tumors: n=4. Lower panel. IHC for the cancer-associated fibroblast marker Fibroblasts Activation Protein (FAP) on ortho_1 Control and sgTWIST1 tumors. Representative images of FAP positive cells characterized by spindle or fusiform morphologies and haphazardly arranged are shown (400x, scale bar: 20 μm). **(C)** Analysis of the protein-protein interactions between the TWIST1-tumor stroma signature (n= 77 genes) and the DE murine stromal genes (n=89). Direct (physical) as well as indirect (functional) interactions analyzed using the String website. All the basic and advanced default settings have been kept but the minimum required interaction score, that has been changed in high confidence (0.7); and the network display options, hiding the disconnected nodes in the network. PPI enrichment *p* value: <1.0^e-16^. Murine stromal genes clustering with the TWIST1 tumor-stroma signature are underlined in black.

Besides, among the up-regulated genes, we noticed the Macrophage Receptor with Collagenous Structure (Marco), which defines a subtype of alternatively-activated M2 tumorassociated macrophages (TAMs) with immunosuppressive functions and involved in tumor progression [33]. Six up-regulated genes of the myofibroblastic signature (*Pvalb*, *Neb*, *Acta1*, *Ttn*, *Myh1*, *Msln*) (Supplementary Fig. S8A) and *Marco* were confirmed by RT-qPCR. For the selected genes of the signature, a reduction in their RNA expression levels was observed in both ortho_sgTWIST1 tumor stroma only, and were undetectable in the tissues from the sc tumors (**Fig. 8B**, Supplementary Fig. S8B). The reduced RNA expression level of *Marco* in sgTWIST1 tumor stroma was validated in all the three *in vivo* models (**Fig. 8B**, Supplementary Fig. S8B). Finally, qualitative validation by IHC with the CAF marker fibroblast-activation protein (Fap) confirmed the presence of CAFs in both Control and sgTWIST1 ortho_1 tumors (**Fig. 8B**).

To analyze the potential interactions existing between the TWIST1-associated tumor-stroma signature and the DE stromal genes, a protein-protein interacting (PPI) network was constructed using the STRING website (https://string-db.org/).The two groups of DE genes clustered separately and had a high level of linkage both among genes of each category and reciprocally (**Fig. 8C**). Two stromal genes reported as myofibroblastic markers, *Acta1*, belonging to the actin family and *Actn2*, a member of the spectrin superfamily, were strongly linked to the network of myofibroblastic genes and connected with the tumor gene cluster, via *TGFB1*, *TGFB3*, *HGF*, *LAMC3* and *LAMA5*, *FIGF* and *HSPB1* (29,30).

## Discussion

In this study, we discovered a role for the embryonic TFs TWIST1 and TWIST2 as prognostic factors in NB. We could reveal the contribution of TWIST1 in enhancing primary and secondary tumor growth and in mediating an aggressive phenotype in *in vivo* NB xenografts.

Furthermore, we identified a TWIST1-associated transcriptional signature, which correlated with outcomes in human primary tumors and activated the TME in an orthotopically-derived xenograft murine model.

TWIST1 and TWIST2 have previously been described as playing a distinct role during embryonic development and having anti-correlated transcriptional expression patterns in spontaneous focal mammary tumors in mice and in human melanoma, colon, kidney, lung and breast cancer [34]. In this study, we show their opposite expression pattern in primary NB and their antithetical prognostic value, highlighting that theTWIST1 expression was correlated with unfavorable NB prognostic factors, metastasis, disease progression, and poor survival. These findings are in line with prior studies conducted on non-pediatric cancers showing the overexpression of TWIST1 in high grade and invasive/aggressive breast, bladder, cervical, ovarian and hepatocellular cancers where it might also serve as prognostic factor for poor outcome [35]. Moreover, we confirmed on larger cohorts of patients previous data showing the association of TWIST1 with MNA NB [14, 15]. Furthermore, TWIST2 was mainly detected in normal tissues and in NB with better prognosis, differently from what observedin several non-pediatric cancers where the upregulation of TWIST2 was associated with a more aggressive phenotype [36–39]. Importantly, we identified TWIST1 as a valid candidate in predicting a poor outcome of patients with LR or no-MNA NB, likewise the HR classification or MNA.

Our *in vivo* investigations on the biological effects of TWIST1 reveal that its loss delays the primary tumor initiation and growth of NB, regardless of the number of cells and the injection site. These data are aligned with prior evidence showing that the suppression of TWIST1 hampers the growth of primary skin papilloma induced by carcinogens [40]; and that the pharmacological inhibition of the Twist-BRD4-Wnt5a signaling axis results in the reduction of tumorigenicity of basal-like breast cancer [41]. Moreover, the overexpression of TWIST1 accelerates tumor establishment and growth of MCF-7-derived breast cancer and transforms mouse embryonic fibroblasts in cells with high tumorigenic potential [34, 42]. In contrast with these findings, TWIST1 was shown as nonessential for primary tumor initiation and growth in several *in vivo* murine models for breast cancer, pancreatic ductal adenocarcinoma and hepatocellular carcinoma, although it seems to play a pivotal role in driving cells migration and invasion [13, 43, 44]. Taken together, these antithetical findings suggest that the role of TWIST1 in carcinogenesis might depend upon the tumor settings as well as on oncogenic drivers.

In our experiments, TWIST1-expressing tumors displayed a phenotype typical of less differentiated NBs. Additionally these tumors were characterized by abundant fascicules of spindle-shaped cells, typical of a mesenchymal-like morphology. The role played by TWIST1 in driving the EMT and in maintaining cells in a mesenchymal state has been widely documented as part of both the morphogenesis during embryonic development, and in the pathogenesis of multiple types of invasive cancers [44–47]. Moreover, several studies demonstrate an association between the EMT and the acquisition of stem-like characteristics in normal and neoplastic epithelial tissues, identifying in TWIST1 the molecular linker between these two biological processes [48–50]. In our study, TWIST1-expressing NB cells were able to grow *in vitro* as neurospheres, known to be enriched in tumor-initiating cells (TIC) exhibiting stem-like features [20]. No differences were observed in the number and in the size of pulmonary micrometastases between the Control and the sgTWIST1 mice. However, TWIST1-expressing NB cells were able to establish pulmonary macrometastases, suggesting an impact of TWIST1 on the last step of the metastatic cascade, the colonization. This process is driven by the self-renewal capability and the proliferative potential of disseminated cancer cells (DCCs) that upon proliferation form macrometastases [51]. Interestingly, in our *in vivo* model both processess were induced by TWIST1. Moreover, we found an increase of TWIST1 in the metastases of NB patients, thus suggesting TWIST1 implication in the formation of clinically detectable metastases.

The contribution of the ECM in the dissemination of cancer cells is well known. Disruption and stiffness of this framework support malignant transformation and cancer progression [26, 52].

In Control tumors expressing TWIST1, we observed a reorganization of the reticulin mesh. Interestingly, a disorganized and cross-linked reticulin network was associated with poor NB prognosis, and a morphometric classification based on variations of both blood vessels and reticulin fibers shape and size was proposed to identify ultra-high risk NB patients [53]. The involvement of TWIST1 transcriptional targets in the degradation/remodeling of the ECM has been demonstrated in both normal embryonic development as well in cancer [26, 45, 54–56]. In our orthotopic model, we found several genes involved in the organization of the ECM and the TME, such as *VCAN*, *ADAMTS19*, *PDGFRA*, *TRIM28* and *PCOLCE*, among the top 20 upregulated by TWIST1, suggesting a role for TWIST1 in defining a permissive microenvironment contributing to the survival and maintenance of cancer stem-like cells. *PCOLCE* is a direct transcriptional target of TWIST1 and is implicated in the regulation of collagen deposition during both early craniofacial development and in osteosarcoma, where it promotes tumor growth, cell migration and invasion [45, 57]. In our study using two cohorts of primary NB, *PCOLCE* was the gene presenting the highest correlation with TWIST1 expression regardless of the amplification status of MYCN, suggesting a role for TWIST1 in the control of *PCOLCE* expression also in primary NB.

For the first time, we identified a NB-associated TWIST1-signature whose elevated expression was found in MNA and HR tumors, and in tumors with a poor survival regardless of the MYCN amplification. In addition, a subgroup of TWIST1-target genes involved in shaping the interface between tumor cells and its stroma was described as TWIST1-tumor-stroma signature. Both signatures were linked to poor survival in primary NB tumors, indicating their biological relevance hence reiforcing the functional role of TWIST1 in NB pathogenesis.

Here we confirm the cooperation between TWIST1 and MYCN in defining a transcriptional program in NB supporting *in vitro* cell proliferation and *in vivo* tumor growth [14, 17]. Moreover, we conclude that these TFs seem to orchestrate distinct functions. Indeed, suppression of TWIST1 in SK-N-Be2c cells and tumors mainly deregulated pathways involved in signaling, nervous system development, migration, adhesion, ECM organization, and cell proliferation.

Interestingly, the genes enriched in the TWIST1-signature are also principally involved in these pathways. On the other side, GO analysis performed on RNAseq data of SK-NBe2c cells downregulated for MYCN through JC1 [17] highlighted a major role for MYCN in controlling the cell cycle regulation and DNA replication. Similar pathways were also identified upon MYCN silencing through JC1 or shRNA in MNA NB cell lines [58], confirming our data.

There are several limitations in our study. First, the use of only one NB cell line to obtain our *in vivo* model could represent an issue in the wider relevance of our findings. Although SK-NBe2c cells are commonly used for NB research, they in fact might not fully represent the biology and diversity of the disease itself. Thus, our observations about the role of TWIST1 in enhancing NB tumor aggressiveness remain to be verified using NB cell lines without MNA as well as primary NB cells. Second, RNAseq analysis was performed on tumors of the ortho_1 experiment, which did not give rise to macroscopic metastases. This was probably caused by extremely rapid tumor growth, which might have prevented the formation of macrometastases. However, this model is suitable for appreciating the effects of TWIST1 on tumor growth capacity and phenotypic features as well as on TME remodeling. Moreover, the main deregulated genes and pathways were consistently altered by TWIST1 between SK-N-Be2c cells and ortho_1 tumors, and the most relevant genes were confirmed in the ortho_2 tumors. Importantly, the biological relevance of the transcriptional program defined by TWIST1 in the SK-N-Be2c ortho_1 xenografts were validated in human primary NB, with the identification of a TWIST1-associated signature and a tumor-stroma signature, both displaying a strong prognostic impact in two cohorts of NB patients. Third, we only focused on the incidence of metastases in the lungs of mice, which occurs in approximately 4% of children with newly diagnosed NB [59]. We did not detect macrometastases in the liver, one of the most frequent sites of infiltration in children together with bone marrow, bone, and lymph nodes. Fourth, the unambiguous identification of the stromal counterpart activated by the tumor-stroma signature remains challenging. Our transcriptomic data suggest an enrichment of M2 TAM and of myofibroblasts, the most abundant stromal cells supporting tumor progression, in TWIST1-positive xenografts. The marked connection observed between the TWIST1-tumor-stroma signature and the stromal DE genes by STRING analysis further support their role in mediating the NB-associated alterations in the tumor stroma. However, the qPCR validation of the stromal genes belonging to the myofibroblastic signature was hampered by sometimes extremely low/undetectable expression levels. This was probably due to the very limited number of stromal cells present in whole tumor lysates. Single cell sequencing could further facilitate the characterization of the impact of TWIST1 on stroma composition. Moreover, precisely identifying CAF by IHC remains difficult due to the lack of specific myofibroblast markers, a common issue in all studies. Finally, it could be argued that an immunocompromised mouse model does not represent the most suitable setup to study TME components. Genetically engineered models spontaneously developing tumors or humanized mouse NB models could represent other valid alternatives to recapitulate the TME composition in NB [60].

In summary and for the first time, our study revealed the prognostic significance of TWIST1 and TWIST2 in NB. The biological impact of TWIST1 on tumor growth and metastatic formation capacity was associated with alterations in the ECM composition and with the establishment of a TME supportive of tumor growth and progression. The transcriptional program activated by TWIST1 in our *in vivo* model of NB further supported these findings and its validation in primary NB unveiled a correlation with HR, progression of the disease and poor prognosis. All our findings strongly indicate a very promising role for targeting TWIST1 in the therapy of HR or relapsed/refractory NB, which remains an almost universally fatal disease.

## Supporting information

This section includes the description of Supplementary Methods; the Supplementary Figures 2 and their Legends; the captions for Supplementary Tables

## Acknowledgements

We thank Dr. Manfredo Quadroni (Protein Analysis Facility, University of Lausanne) and his team for their help with the secretome analysis. Dr Jessica Dessimoz (Histology Core Facility of the EPFL) and Janine Horlbeck (Mouse Pathology Facility) for their help with the IHC and tissue staining. Dr Julien Marquis (Lausanne Genomics Technologies Facility) for his technical advice in the targeted sequencing analysis of the indels in TWIST1 gene. This work was supported by grants from Kinderkrebsforschung Schweiz (to A.M.M.), the Novartis Foundation for Medical-Biological Research (to A.M.M.), the Hubert Gouin Association (to A.M.M.), and the FORCE foundation (to A.M.M. and R.R.).

## Author contributions

M.V.S. performed all major experimental work, with the technical help of K.B.B., M.V.S. and A.M.M. analyzed the data, prepared figures and drafted the manuscript, J.M.J and N.J. performed *in vivo* xenograft implantations, K.N.A. constructed the LentiCRISPR v2-sgTWIST1 vectors, J.Y.S. provided the TMA., H.S. performed the TMA analysis and the interpretation of the related data, A.P. provided help in the TMA analysis, V.P. conducted the bioinformatics analysis, N.R. performed pathological analyses of the xenografts, R.R. interpreted the data and edited the manuscript, A.M.M. designed, supervised the study and coordinated experiments. All authors read, commented and approved the final manuscript.

## Data availability

All data generated during this study are included in this article (and its Supplementary Information file). The RNAseq, proteomics and image corresponding datasets can be accessed at the GEO public repository using the accession number GSE160765; at the Proteomics Identifications Database (PRIDE) using the accession number PXD024200; and at the Zenodo repository with the doi: 10.5281/zenodo.4543478, respectively. The RNAseq data of SK-N-Be2c JC1 samples were obtained from GEO, using the accession number GSE80153. The relevant data that support the findings of this study are available from the corresponding author upon reasonable request. Source data are provided with this paper.

