## Supplementary material for "TWIST1 expression is associated with high-risk Neuroblastoma and promotes Primary and Metastatic Tumor Growth": This section includes the description of Supplementary Methods; the Supplementary Figures 2 and their Legends; the captions for Supplementary Tables

### Supplementary Information:

This section includes the description of Supplementary Methods; the Supplementary Figures and their Legends; the captions for Supplementary Tables S1 to S14.

#### Supplementary Methods:

##### Cancer Cell Line Encyclopaedia (CCLE) database analysis

The CCLE database (https://portals.broadinstitute.org/ccle) was used to compare the mRNA expression level of TWIST1 and TWIST2 in a range of 36 different cancer types (Affymetrix arrays: Affimetrix Human Genome U133, Plus 2.0). Quality filtering and normalization were performed using Robust Multi-array Average (RMA) and quantile normalization.

##### TWIST1 knock out through CRISPR/Cas9 technology

Two sgRNAs targeting the first exon of the TWIST1 gene were chosen in the published sgRNA library[1]. Oligos were designed as follow: sgTWIST1-1: forward 5'-CACCGCGGGAGTCCGCAGTCTTACG-3'; reverse 5'-AAACCGTAAGACTGCGGACTCCCGC-3'; sgTWIST1-2: forward 5'-CACCGCTGTCGTCGGCCGGCGAGAC-3'; reverse 5'-AAACGTCTCGCCGGCCGACGACAGC-3'. The lentiviral vector lentiCRISPR v2 [2] was obtained from Adgene (Cambridge, USA). LentiCRISPR v2-sgTWIST1 plasmids were constructed according to the manufacturer's instructions (Adgene), and used to transduce Control cells. Virus production and lentiviral infections were performed as previously described[3]. Transduced SK-N-Be2c, LAN-1 and NB1-M cells were selected 24 h post-infection with either 5 µg/ml for SK-N-Be2c or 1 µg/ml for LAN-1 and NB1-M cells of puromycin (Gibco). Clones were isolated by limiting dilution cloning in a 96-wells plate from the Control and sgTWIST1 #1 of LAN-1 and SK-N-Be2c cell lines. Only clones derived from a single colony were selected and further screened by Immunoblotting. To avoid potential problems caused by variability during single-cell clonal expansion, we pooled 5 and 7 clones for the Control and sgTWIST1 SK-N-Be2c cells, respectively; for the LAN-1 cells, 6 clones were pooled for both

Control and sgTWIST1 cells. SK-N-Be2c-Control and sgTWIST1 pools of cells were further transduced with a lentiviral vector expressing the mCherry gene (Ex-NEG-Lv244 from GeneCopoeia$^{TM}$). These cells expressing the mCherry gene were used for the ortho_2 and the sc xenograft experiments. In addition, genome editing in SK-N-Be2c sgTWIST1 cells was verified by NGS sequencing. Briefly, PCR amplicons were designed across the TWIST1 genomic regions targeted by the sgRNAs to examine generation of indels. A first PCR of 20 cycles was performed using the primers Twist1-nest-F: 5'-GCAAGAAGTCTGCGGGCTGTGG-3' and Twist1-nest-R: 5'-GGATGATCTTCCGCAGCGCG-3', followed by purification with QIAquick PCR purification kit (QIAGEN). Then we run a second PCR of 10 cycles with nested primers containing Illumina overhang adapter sequences: Illumina-P5-Twist1-F: 5'-TCGTCGGCAGCGTCAGATGTGTATAAGAGACAG AAGAAGTCTGCGGGCTGTGGCG-3'; Illumina P7-Twist1-R: 5'-GTCTCGTGGGCTCGGAGATGTGTATAAGAGACAGCGCTCCCGCACGTTGGCCATG-3'. Both PCR were performed using the 2xKAPA HiFi HotStart ReadyMix as follow: 95°C for 5 minutes, 10 cycles at 98°C for 20 s, 73°C for 30 s, 72°C for 30 s; then 72°C for 5 min. Finally, a third PCR was performed to attach Illumina adaptors and barcodes to samples according to manufacturer's instructions. Amplicons were purified by using AMPure XP (Beckman Coulter, Indianapolis, USA), and sequenced with a MiSeq Micro 300 (Illumina Inc., San Diego, USA) at the Lausanne Genomics Technologies Facility (GTF) (https://wp.unil.ch/gtf). After high throughput sequencing reads, PCR amplicons were checked for indels using CRISPResso (https://crispresso.pinellolab.partners.org). The results of the sequencing of the bulk population of SK-N-Be2c-sgTWIST1 cells, the 7 derived clones, and the 4 sgTWIST1 ortho_1 tumors are shown in Supplementary Table S13. Surprisingly, in all the sgTWIST1 clones we detected the same three main indels. Note that three alleles found in the SK-N-Be2c cells, indicating a triploidy of this genomic region as previously described [4].

**Neurosphere Assay**

NB cells (2x$10^4$ cells/ml) were cultured plated in duplicate in Neural Crest Stem Cell culture medium (NCSC) [DMEM/F12 (Gibco) supplemented with 1 % penicillin/streptomycin (Gibco), 2 % B27 (Gibco), 20 ng/ml human recombinant bFGF (Peprotech, Rocky Hill, USA), 1% N2 (Gibco), 2-Mercaptoethanol 50µM (AppliChem, Darmstadt, Germany) 15% Chicken Embryo Extract, 20 ng/ml IGF-1 (Peprotech), Retinoic Acid 110 nM], using poly-Hema-coated six wells plates to prevent cell adhesion. After 7 days in culture, pictures of sphere were taken using an Olympus IX53 inverted microscope (Olympus, Shinjuku, Japan) and acquired with the Olympus cellSense imaging software. Spheres were dissociated with StemPro Accutase Cell Dissociation Reagent (Gibco, A11105-01) and the number of cells recovered after the dissociation of spheres was determined using the trypan blue exclusion method.

**Proliferation assay**

Briefly, 1.2*$10^4$ MNA (SK-N-Be2c and LAN-1) and 3*$10^4$ no-MNA (NB1-M) cells were seeded in quadruplicate in 96-wells plate in DMEM/FCS. Cell proliferation was assessed using the CellTiter 96® Aqueous Non-radioactive Cell proliferation Assay (Promega, Madison, WI, USA) according to the manufacturer's protocol.

**Real-Time qPCR**

cDNA were prepared from 0.5 or 1 µg of RNA for the validation of human or murine genes, respectively using PrimeScriptTM reagent kit according to the manufacturer's instruction (TAKARA Bio Inc., St.Germain-en-Laye, France). The expression level of selected genes was validated by semi-quantitative real-time PCR in duplicates using primer pairs described in Supplementary Table S14 and the QuantiFast SYBR® green kit (Qiagen, Hilden, Germany). Cycling conditions were: 5 min at 95°C, 40 cycles of 10 sec at 95°C, 30 sec at 60°C, and 1 sec at 72°C with the Rotor Gene 6000 real-time cycler (Corbett, Qiagen). Gene expression levels were determined by normalization to either HPRT1 (human genes) or β-actin (murine genes) housekeeping genes using the ΔCt method.

**Immunohistochemistry**

Histopathological analyses were performed on blocks embedded in paraffin at the Mouse Pathology Facility of Lausanne University (Epalinges, Switzerland) and at the Histology Core Facility of the EPFL (Lausanne, Switzerland). Thin tumor sections (3 µm thick) were de-waxed and rehydrated and then stained with hematoxylin and eosin; Gomori's for reticulin or type III collagen detection, or IHC was performed using primary antibodies (Supplementary Table S13). Staining were imaged using an Olympus BX43 light microscope and then acquired with the Olympus cellSense imaging software or using the EVOS5000 imaging system (Life Technologies); or whole slides were scanned at the 20x magnification using the NanoZoomer S60 Digital slide scanner C13210 (Hamamatsu, Japan) for the VCAN; and the Zeiss Axioscan Z.1 for the Alu. Visualization and analyses of the scans was performed using the QuPath imaging software (https://qupath.github.io). For the quantification of VCAN and Alu positive probe II, the entire surface either of the tumor or of three subsequent lung sections separated by a depth of 100 µm has been taken into consideration, respectively. Quantification was perfomed with the QuPath software according to the following pipeline: for the VCAN, select a region of the tumor (magnification: 12.48X), analyze, cell analysis, positive cell detection; for the Alu positive probe II, a polygonal region has been traced around a group of positive cells and the area has been quantified automatically. Options modified from the default parameters for the VCAN quantification: image type: Brightfield (other); detection image: optical density sum; intensity threshold parameters: threshold 1+: 0.3.

**Immunoblotting**

NB cells and SK-N-Be2c-derived ortho tumors were lysed in NP40 buffer (50 mM Tris-HCl pH 8.0; 150 mM NaCl; 1% NP-40 and 1x protease inhibitor cocktail (Complete mini, EDTA-free, Roche, Mannheim, Germany). Cell lysates were centrifuged at 15'000 rpm at 4°C for 10 min. Supernatants were recovered and protein concentrations were measured using Bradford method (Bio-Rad Laboratories, Hercules, CA, USA). Equal amount of total protein lysate (40 µg for DE genes validation; 50 µg for TWIST1 and MYCN validation) was loaded onto 4-15%

precast polyacrylamide Mini-PROTEAN TGX gel (Bio-Rad Laboratories). Proteins were transferred to PVDF membrane (Immuno Blot,Bio-Rad Laboratories) and blocked with 3% nonfat dry milk in TBS-T 0.1%, incubated overnight at 4˚C with primary antibodies (Supplementary Table S12) and then for one hour with the appropriate secondary antibodies (Supplementary Table S12). Proteins were imaged using either the WesternBright Sirius Kit or the WesternBright Quantum Kit (Advansta Inc., San Jose, CA, USA) and the Fusion FX Spectra multimodal imaging platform (Vilber Lourmat, Marne-la-Vallée, France). Quantification of immunoreactive bands was performed using the ImageJ software according to the following pipeline: analyze, gel, select 1 lane, plot lanes, manually define the area corresponding to each band, wand tool to quantify each selected area.

### Supplementary Figures

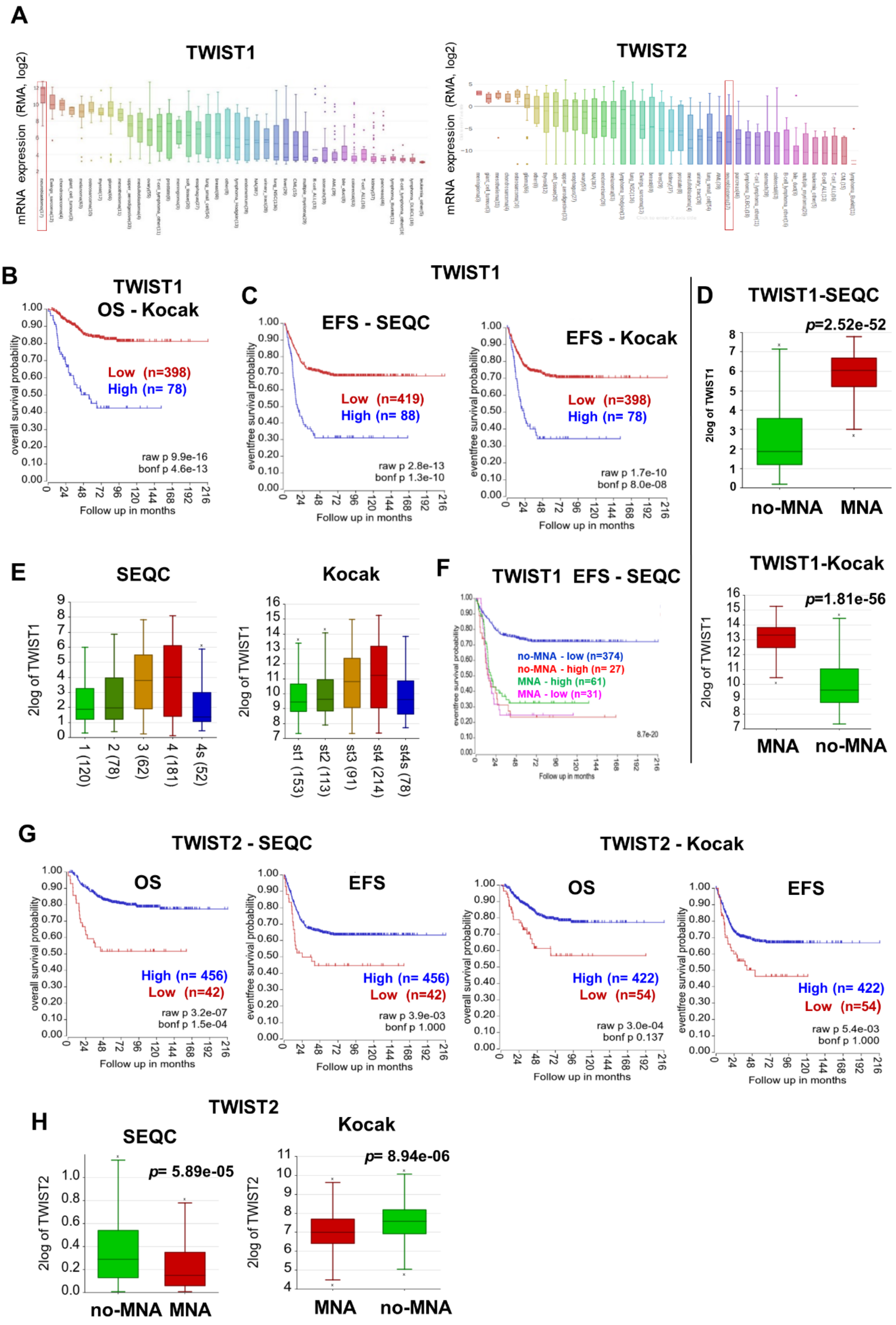

**Supplementary Fig. S1**. **TWIST1 and TWITS2 RNA expression in NB cells and tumors**. (**A**) Box plot showing the mRNA expression levels of TWIST1 (left) and TWIST2 (right) in a panel of 40 cancer cell lines in the CCLE database. The numbers in the brackets correspond to the numbers of cell lines per tumor types. (**B**) Kaplan-Meier OS curve associated with TWIST1 expression in the Kocak dataset of primary NB tumors (expression cutoff = 6701.9) (n=476 with survival data). (**C**) EFS associated with TWIST1 expression in the SEQC (left panel, expression cutoff: 44.441) and Kocak (right panel, expression cutoff = 6701.9) datasets. (**D** and **E**) Box-and-whisker plots of TWIST1 expression in MNA and no-MNA tumors (**D**); and in tumors with distinct INSS stages in the indicated datasets (**E**). (**F**) Kaplan-Meier EFS curves showing the stratification of patients of the SEQC dataset according to MYCN status and TWIST1 expression (high or low). (**G**) OS and EFS according to TWIST2 expression in the SEQC (left panels, expression cutoff: 1010) and Kocak (right panels, expression cutoff: 86.0) datasets. (**H**) Box-and-whisker plots of TWIST2 expression in no-MNA and MNA tumors in the SEQC and Kocak datasets.

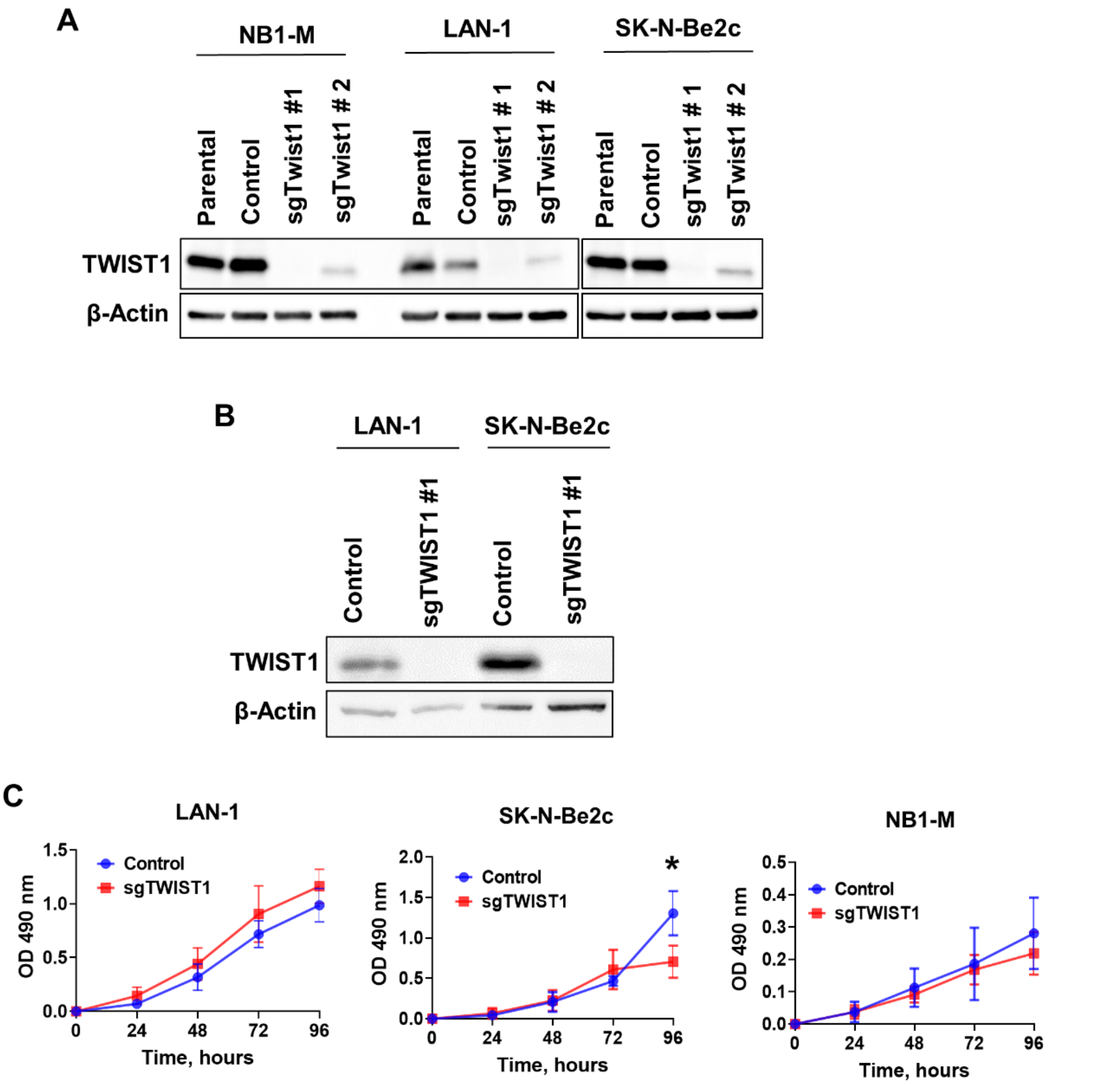


**Supplementary Fig. S2. Validation of the CRISPR/Cas9-mediated TWIST1 KO and its impact on NB cell proliferation *in vitro*.** (**A**) Immunoblotting for the detection of TWIST1 protein expression and β-actin (as the loading control) in the bulk populations of NB cells before (Parental) and after the lentiviral infection: Control vector, sgTWIST1#1 (sgTWIST1, from now on) and sgTWIST1#2. (**B**) TWIST1 protein expression in the LAN-1 and SK-N-Be2c pool of clones for Control and sgTWST1 (see Material&Methods). (**C**) Cell proliferation of Control and sgTWST1 NB cell lines measured by MTS/PMS assay from 24h to 96h. Mean OD 490nm ± SD of three (SK-N-Be2c) and four (LAN-1 and NB1-M) independent experiments performed in quadruplicates are shown. Statistical analysis was done using the Holm-Sidak multiple t-test (α=0.05), without assuming a consistent SD. *$p$= 0.0376 in SK-N-Be2c.

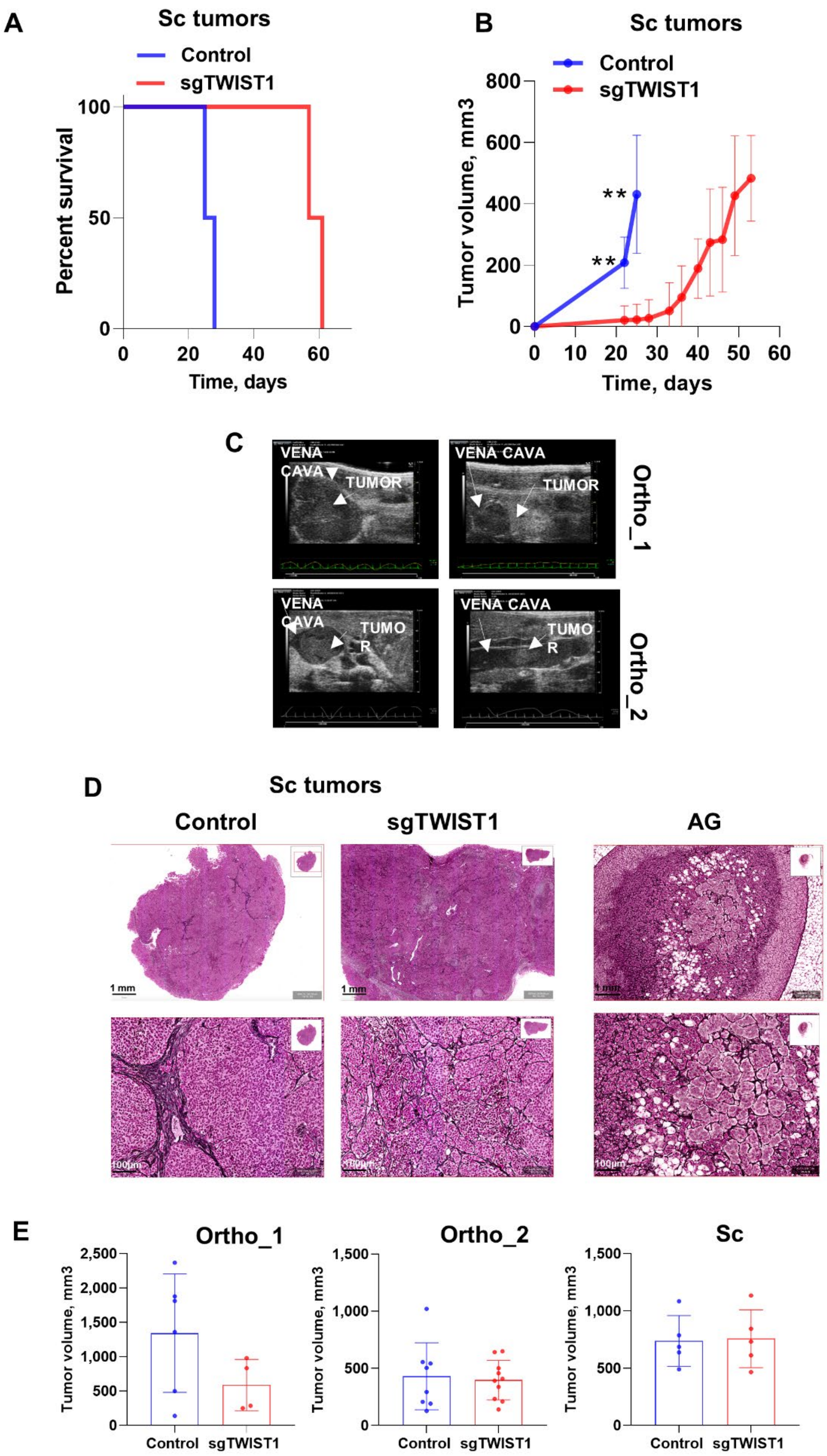
A
Sc tumors
Control
sgTWIST1
Percent survival
Time, days
B
Sc tumors
Control
sgTWIST1
Tumor volume, mm3
Time, days
C
VENA CAVA
TUMOR
Ortho_1
Ortho_2
D
Sc tumors
Control
sgTWIST1
AG
1 mm
100µm
E
Ortho_1
Ortho_2
Sc
Tumor volume, mm3
Control
sgTWIST1

**Supplementary Fig. S3. TWIST1 KO diminishes the tumor growth and invasive capacities of SK-N-Be2c cells and affects the Collagen III/reticulin network organization.** (**A**) Kaplan-Meier survival curve of mice implanted subcutaneously with SK-N-Be2c-Control or -sgTWIST1 cells. Mice were sacrificed once tumors reached approximately 700 $mm^3$. Tumor take: 100% (5/5) in both groups. Median survival in the Control vs sgTWIST1 groups: 26.5 vs 59 days. Gehan-Breslow-Wilcoxon test: ****p*=0.0002. (**B**) Tumor growth curve for the sc experiment. Data are plotted as the mean tumor volume ± SD. Mann-Whitney t-test: ***p*= 0.0079 at both 22 and 25 days. (**C**) Representative ultrasound images of tumor cell intravasation in the vena cava of 2 distinct ortho_1 and ortho_2 Control mice. (**D**) Representative images of Gomori's staining showing the ECM architecture of Control and sgTWIST1 sc tumors (scale bars: top 1mm; bottom 100µm) as compared to the normal AG (scale bars: top 200 µm; bottom 100 µm). (**E**) Graphs illustrating the mean tumor volumes at sacrifice ± SD. Ortho_1: mean Control= 1343 $mm^3$, n=6; sgTWIST1= 587 $mm^3$, n=4; Mann-Whitney test: *p*= 0.257. Ortho_2*:* mean Control= 430 $mm^3$, n= 8; sgTWIST1= 397 $mm^3$, n=10; unpaired t-test: *p*= 0.. *SC*: mean Control: 737 $mm^3$, n=5; sgTWIST1= 757 $mm^3$, n=5; Mann-Whitney: *p*>0.999.

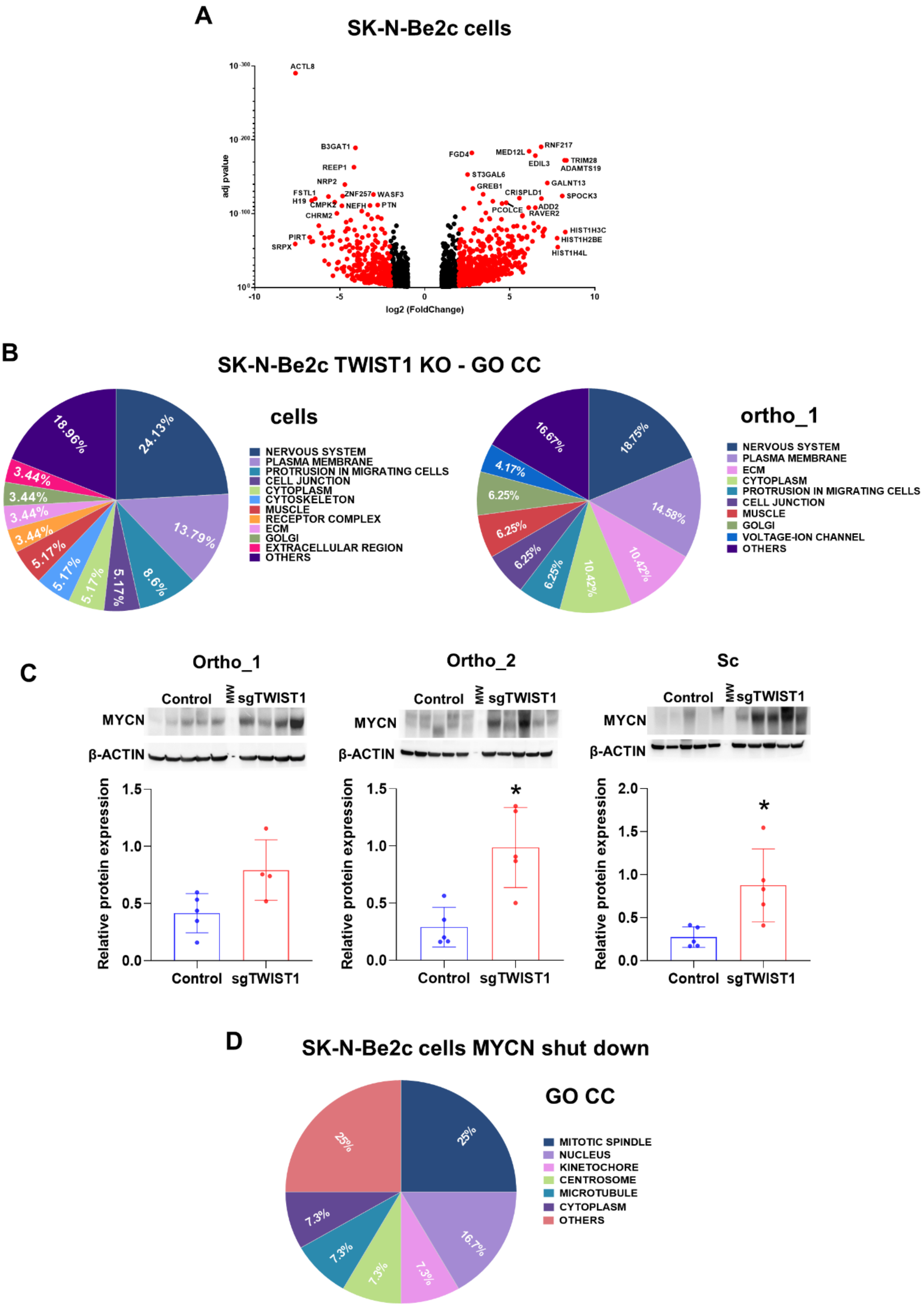
A
SK-N-Be2c cells
adj pvalue
log2 (FoldChange)
ACTL8
B3GAT1
REEP1
NRP2
FSTL1
H19
CMPK2
ZNF257
WASF3
NEFH
PTN
CHRM2
PIRT
SRPX
FGD4
MED12L
RNF217
EDIL3
TRIM28
ADAMTS19
ST3GAL6
GREB1
GALNT13
CRISPLD1
SPOCK3
PCOLCE
ADD2
RAVER2
HIST1H3C
HIST1H2BE
HIST1H4L
B
SK-N-Be2c TWIST1 KO - GO CC
cells
NERVOUS SYSTEM
PLASMA MEMBRANE
PROTRUSION IN MIGRATING CELLS
CELL JUNCTION
CYTOPLASM
CYTOSKELETON
MUSCLE
RECEPTOR COMPLEX
ECM
GOLGI
EXTRACELLULAR REGION
OTHERS
24.13%
13.79%
8.6%
5.17%
5.17%
5.17%
5.17%
3.44%
3.44%
3.44%
3.44%
18.96%
ortho_1
NERVOUS SYSTEM
PLASMA MEMBRANE
ECM
CYTOPLASM
PROTRUSION IN MIGRATING CELLS
CELL JUNCTION
MUSCLE
GOLGI
VOLTAGE-ION CHANNEL
OTHERS
18.75%
14.58%
10.42%
10.42%
6.25%
6.25%
6.25%
6.25%
4.17%
16.67%
C
Ortho_1
Ortho_2
Sc
Control
MW
sgTWIST1
MYCN
β-ACTIN
Relative protein expression
Control
sgTWIST1
*
D
SK-N-Be2c cells MYCN shut down
GO CC
MITOTIC SPINDLE
NUCLEUS
KINETOCHORE
CENTROSOME
MICROTUBULE
CYTOPLASM
OTHERS
25%
16.7%
7.3%
7.3%
7.3%
7.3%
25%

**Supplementary Fig. S4. Distinct transcriptional programs are affected upon TWIST1 KO and MYCN shut down.** (**A**) Volcano plots showing the distribution of the gene expression fold changes and adjusted *p* value for the DE genes in SK-N-Be2c-Control versus SK-N-Be2c–sgTWIST1 cells. Genes with False Discovery Rate (FDR) $< 0.05$ and absolute value (av) of $\log2(FC) \geq 1$ were considered as DE; in red genes with av of $\log2(FC) \geq 2$, in black genes with av of $\log2(FC) \geq 1$ and $<2$. Positive and negative x-values represent genes either up or down-regulated by TWIST1 respectively. (**B**)**.** Illustration of the cellular components gene sets found enriched by GO analyses (GO CC) in the DE genes following TWIST1 KO for both SK-N-Be2c cells (left panel) and ortho_1 tumors (right panel). Data are reported as the repartition (in %) of the diverse pathways identified with a FDR $< 0.01$ (n=58 for cells, n=48 for tumors). (**C**) Immunoblotting for MYCN protein and $\beta$-actin as control (upper panel) and densitometric quantification of MYCN expression relative to $\beta$-ACTIN (lower panel) in SK-N-Be2c-derived tumors of the 3 *in vivo* experiments. Expressions relative to $\beta$-ACTIN were plotted as individual data with mean ± SD. Mann Whitney test. **p*= 0.0159 in ortho_2 and sc. Ortho_1: n= 5 Control; n= 4 sgTWIST1; ortho_2 and sc: n= 5 Control; n= 5 sgTWIST1. (**D**) Illustration of the GO CC gene sets found enriched in the DE genes of SK-N-Be2c cells upon JC1-medited MYCN shutdown. RNAseq data of SK-N-Be2c cells treated with JC1are during 24h or DMSO as control were uploaded (GSE80154, see Methods) (Zeid et al.). Genes with False Discovery Rate (FDR) $< 0.05$ and absolute value (av) of $\log2(FC) \geq 1$ were considered as DE. Data are reported as the repartition (in %) of the diverse pathways identified with a FDR $< 0.01$ (n=24).

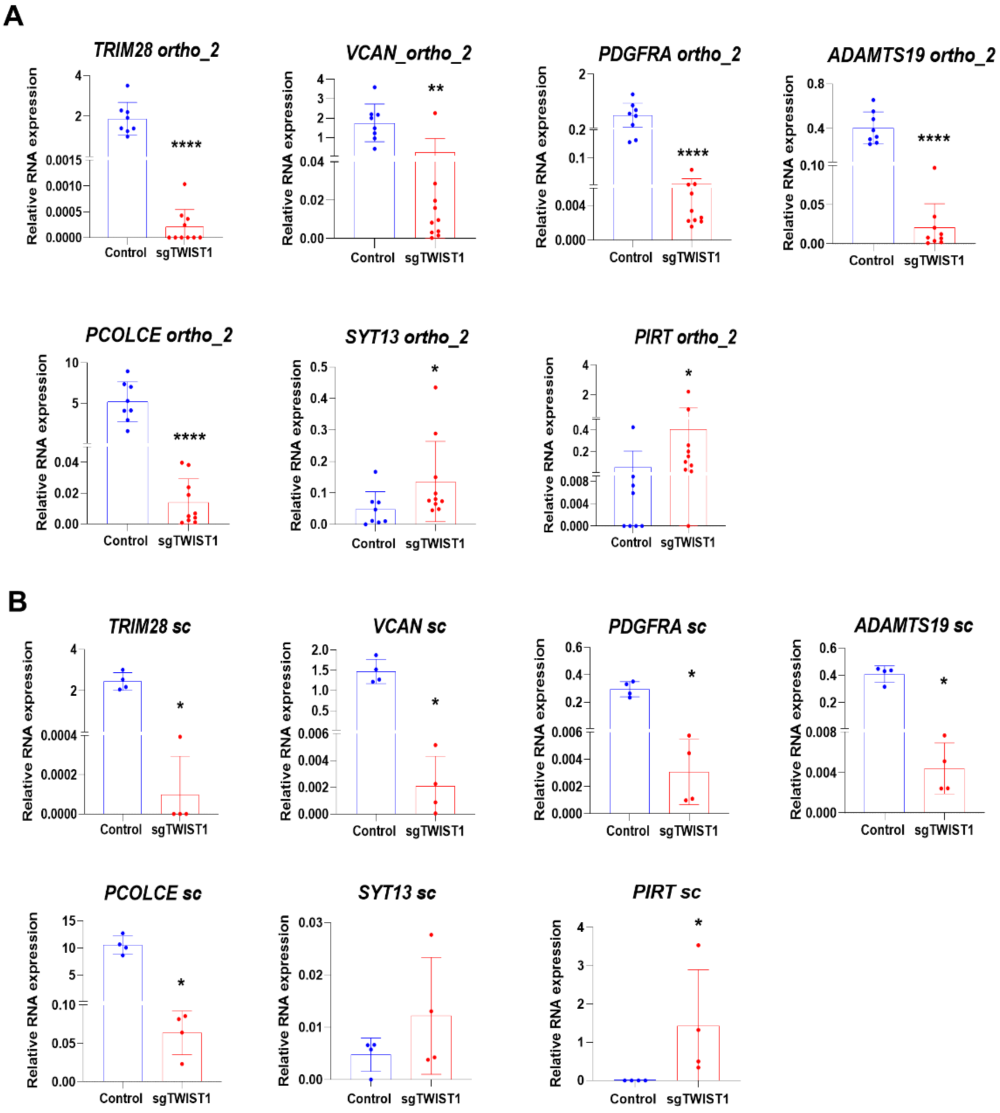


**Supplementary Fig. S5. Validation of TWIST1-mediated deregulation of selected TWIST1 target genes at the RNA level in SK-N-Be2c-derived tumors**. **A** and **B**: mRNA expression levels of the selected TWIST1 target genes relative to *HPRT1* analyzed by RT-qPCR are plotted as individual values with mean ± SD for the indicated *in vivo* experiments**.** Numbers of tumors analyzed for ortho_2 (**A**): Control n= 8, sgTWIST1 n= 10; and for sc (**B**): Control n= 4, sgTWIST1 n= 4. Statistical analysis was performed using Mann Whitney test for all genes but *PCOLCE in* ortho_2 experiment (unpaired t-test: ****$p$<0.0001). Ortho_2: *TRIM28*: ****$p$=0.0001 2; *VCAN*: **$p$=0.0021; *PDGFRA* and *ADAMTS19*: ****$p$=0.0001; *PIRT*: *$p$=0.0146; *SYT13*: *$p$=0.0266; sc: *TRIM28*: *$p$=0.; *VCAN*, *PDGFRA*, *ADAMTS19* and *PIRT*: *$p$=0.0286; SYT13: *$p$=0.0266.

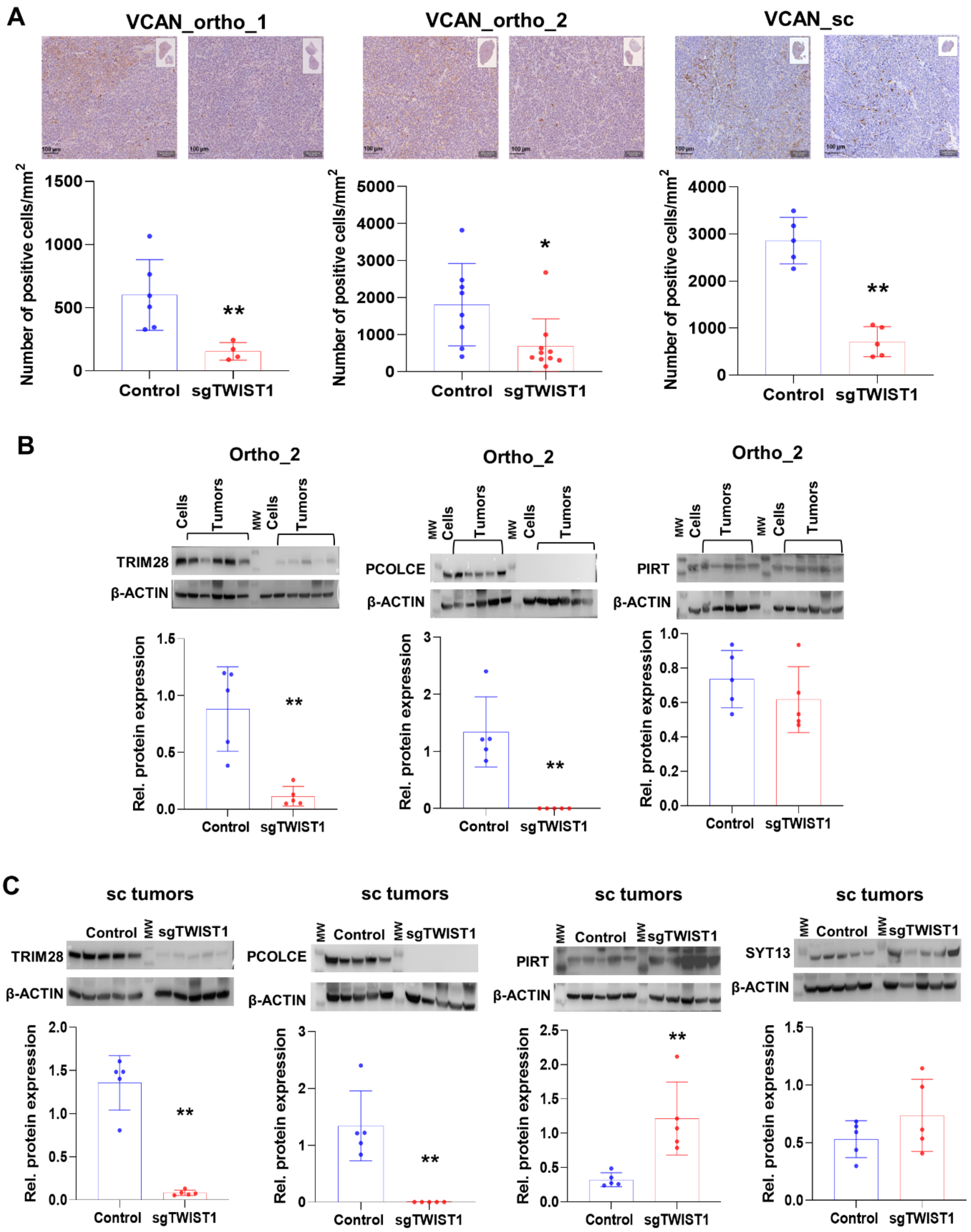


**Supplementary Fig. S6. Analysis of the protein expression levels of selected TWIST1 target genes in SK-N-Be2c-derived tumors.** (**A**) Upper panels: representative images of IHC for VCAN in the indicated tumors (scale bar =100 µm); lower panels: quantification of VCAN staining (Qpath software on total area of each section). Mann-Whitney: ortho_1:***p*=0.0095, n=6 Control and n=4 sgTWIST1; ortho_2: **p*= 0.0155, n=8 Control and n=10 sgTWIST1; sc: ***p*=0.0079, n=5 Control and n=5 sgTWIST1. (**B** and **C**). Relative protein expression as determined by immunoblotting for the selected genes in the ortho_2 (**B**) and the sc (**C**) tumors. Upper panel: Representative images of immunoblotting for TRIM28, PCOLCE, PIRT and SYT13 (β-ACTIN as the control); MW: molecular weight; lower panel: densitometric quantifications of immunoreactive band densities. Expressions relative to β-ACTIN were plotted as individual data with mean ± SD. Mann Whitney test: ***p*=0.0079 for all comparisons. Ortho_2 and sc: n=5 Control and n=5 sgTWIST1.

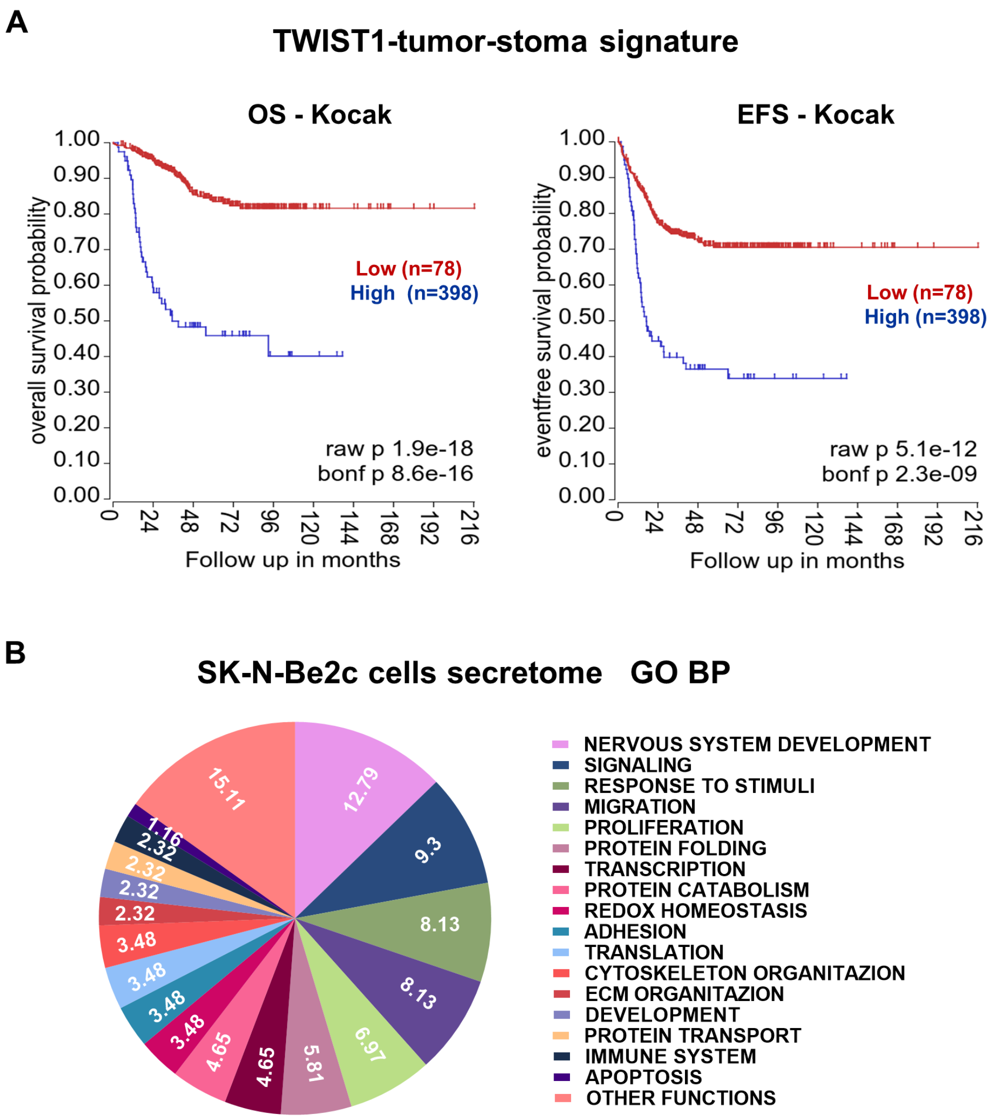


**Supplementary Fig. S7. Correlation between the TWIST1-tumor-stroma signature in the Kocak NB dataset and the outcome of patients.** (**A**) Kaplan-Meier plots showing the correlation between a high level of the paracrine signature expression and a poor OS and EFS of NB patients in the Kocak dataset. Expression cutoff for both curves: 0.14. (**B**) Illustration of biological processes (BP) found enriched by gene ontology analysis for the DE proteins in SK-N-Be2c cell secretome. Data are reported as the repartition (in %) of the diverse BP identified with a FDR < 0.01 (n=50).

**Figure S7**

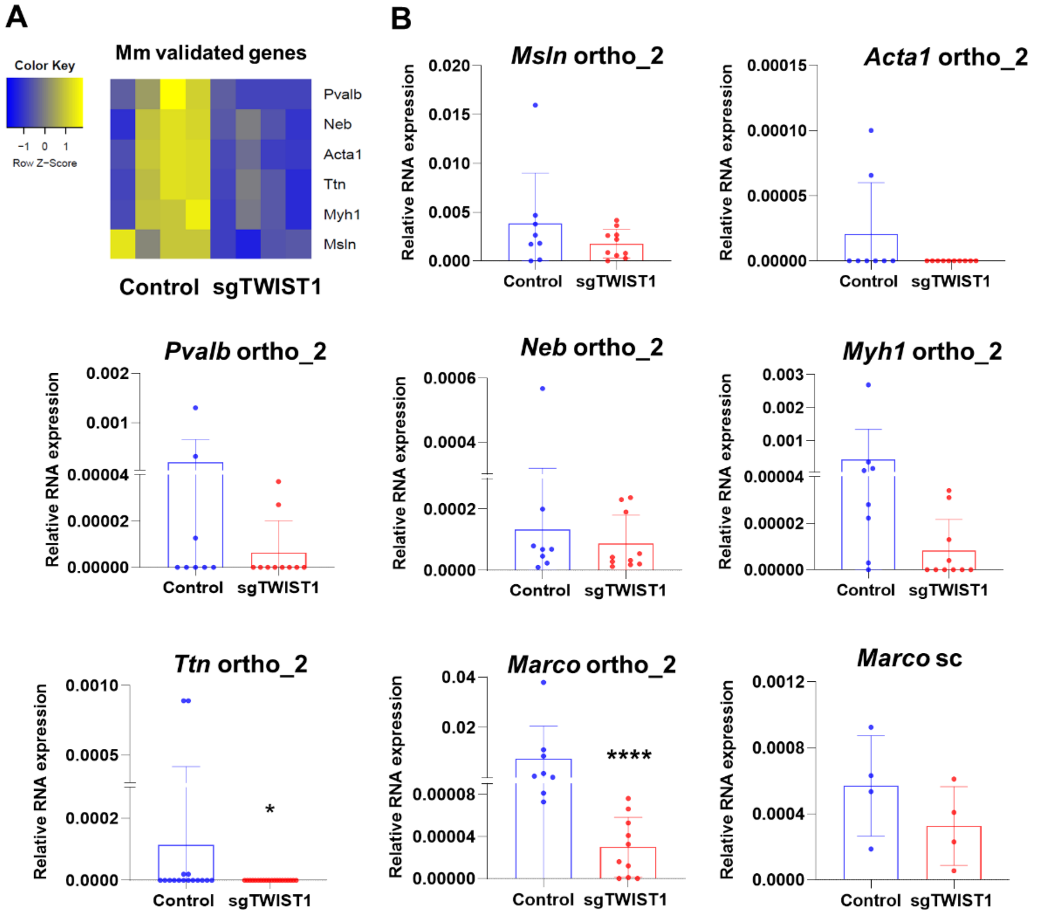


**Supplementary Fig. S8**. **Validation of selected genes of the myofibroblast signature in the ortho_2 and sc tumors by real-time PCR.** (**A**) Heatmap showing the RNA expression levels (z-score) of the Myofibroblasts signature selected genes as determined by RNAseq analysis in ortho_1 tumors. (**B**) mRNA expression levels for the selected myofibroblast genes and *Marco* relative to *β-actin* as determined by RT-qPCR. Data are plotted as individual values with mean ± SD. Mann Whitney test: Ortho_2: *Ttn*: **p*= 0.0309; *Marco*: *****p*<0.0001. Ortho_2 tumors: Control n=8; sgTWIST1 n=10; sc tumors: Control n=4, sgTWIST1 n=4.

### Supplementary Tables

**Captions for Supplementary Table S1 to Table S14:**

Table S1: Composition of the TMA and expression of TWIST1 and TWIST2 in NB primary tumors, metastases and control tissues.

Table S2: List of raw counts, RPKM and differentially expressed (DE) human protein coding genes in SK-N-Be2c-Control and -sgTWIST1 Cells and Tumors.

Table S3: List of pathways identified upon TWIST1 KO by gene ontology analysis in SK-N-Be2c Cells and Tumors.

Table S4: List of raw counts and DE genes after MYCN shutdown with JC1 for 24h.

Table S5: List of pathways identified by gene ontology analysis deregulated by MYCN in SK-N-Be2c cells.

Table S6: Gene lists for the identification of the TWIST1-signature.

Table S7: List of genes of the TWIST1-tumor-stroma signature.

Table S8: Composition of the SK-N-Be2c cell secretome.

Table S9: List of pathways identified by gene ontology analysis on the TWIST1-deregulated proteins of the secretome.

Table S10: List of all murine protein coding genes detected by RNAseq in SK-N-Be2c-Control and -sgTWIST1 ortho_1 tumors and the 89 DE genes.

Table S11: Summary of myofibroblast markers and principal muscle structure-specific genes.

Table S12: Gene ontology analysis on murine changing genes upon TWIST1 KO.

Table S13: Illustration of the insertions/deletions generated in the TWIST1 gene.

Table S14: List of primers and antibodies used in the study.
